# Jack of all trades: genome assembly of Wild Jack and comparative genomics of Artocarpus

**DOI:** 10.1101/2022.09.02.506353

**Authors:** Ajinkya Bharatraj Patil, Sai Samhitha Vajja, S. Raghavendra, B.N. Satish, C.G. Kushalappa, Nagarjun Vijay

## Abstract

Known as breadfruits for their diverse nutritious fruits, the Artocarpus (Moraceae) genus is prized for its high-quality timber, medicinal value, and economic importance. Breadfruits are native to Southeast Asia but have been introduced to other continents. The most commonly cultivated species are *Artocarpus heterophyllus* (Jackfruit) and *Artocarpus altilis* (Breadfruit). With numerous smaller but nutritionally comparable fruits on a large tree, *Artocarpus hirsutus*, also called “Wild jack” or “Ayani” is an elusive forest species endemic to Indian Western Ghats. In this study, we sequenced and assembled the whole genome of *Artocarpus hirsutus* sampled from sacred groves of Coorg, India. We compared our Wild jack genome with previously published Jackfruit and Breadfruit genomes to decipher demography and evolution. Demographic history reconstruction indicates a stronger effect of habitat rather than phylogeny on the population histories of these plants. Repetitive genomic regions, especially LTR Copia, strongly affected the demographic trajectory of *A. heterophyllus*. Upon further investigation, we found a recent lineage-specific accumulation of LTR Copia in *A. heterophyllus*, which results in its larger genome size. Gene family evolution and signatures of selection identified multiple genes from starch, sucrose metabolism, plant hormone signal transduction, etc., in *Artocarpus* species. Our comparative genomic framework provides important insights by incorporating endemic species such as the Wild jack.

## Introduction

Genus Artocarpus (Moraceae), or “Breadfruits,” are tropical plants famous for their nectary and fleshy fruits (Jarrett, 1977). This genus comprises ~70 species with considerable variability in size, height, flower/fruit morphology, developmental processes, and functional properties (Zerega et al., 2010; Gardner et al., 2021). Almost all of the members of the genus provide a rich resource of food, timber, and other valuable products, popularising them in their native regions (Jagtap and Bapat, 2010; Xavier et al., 2014; Ragone, 2018). Due to these properties, some species have been introduced to various parts of the world. The two most widely distributed domesticated species, *Artocarpus heterophyllus* (Jackfruit) and *Artocarpus altilis* (Breadfruit), currently have oriental distribution in the tropical and subtropical regions (Zerega et al., 2010; Williams et al., 2017). Artocarpus trees are native to the region extending from the Western Ghats, South-East Asia to the Oceanic Islands. A recent study suggested diversification of Artocarpus from Borneo followed by subsequent dispersal and divergence during the Miocene (Williams et al., 2017). But there are multiple fossils found in India, suggesting the occurrence of Artocarpus species during the Palaeocene, much before Miocene (Mehrotra, RC, Prakash, U Bande, 1984; Srivastava, 1998). Moreover, the Bornean origin of Artocarpus suggests overwater or overland dispersal over large distances as the only possibility for Indian Artocarpus species to exist, which feels unlikely (Williams et al., 2017). Hence, the biogeographical history of these plants is yet to be established and is a matter of further research. Differences in bioclimatic properties of their habitats and the fauna involved in its pollination/dispersal might have played an instrumental role in adapting these species by developing divergent characteristics from their ancestral counterparts.

Artocarpus trees are well known for their diversity of unique unisexual inflorescences and composite syncarpous fruits (Jarrett, 1977). The phenotypic diversity among the syncarps is such that the taxonomy of this genus is entirely dependent upon inflorescence morphology and structure (Zerega et al., 2010). Even though the focus has been on the floral diversity for delineating these species, these plants have evolved several other species-specific characters. For instance, the trees of *A. heterophyllus* reach the height of 15-20 meters and have reticulate branching close to the soil (**Figure 1**). The tree of *A. altilis* reaches up to 30 meters and is moderately branched at a medium height from the ground. In contrast to these two, Wild Jack (*Artocarpus hirsutus*) are large forest trees that reach above 50 meters, some extending till 70 m with no branching until the apices. The male inflorescences differ in all three species. *A. heterophyllus* has smaller cylindrical inflorescence than *A. altilis*, which has longer and thicker apices. In contrast, *A. hirsutus* has a thin, long, filamentous stalk as the male inflorescence differs entirely from these two species. Female inflorescences also differ in these three species, so their fruit morphology is quite diverse. Jackfruit (*A. heterophyllus*) bears multiple, low-hanging, larger, ellipsoidal, fleshy, nectary, and green sheathed fruits of sizes up to 100 cm. The Breadfruit (*A. altilis*) bears numerous medium-sized, oval, starchy, and green sheathed fruits of 12-20 cm, hanging at apices of branches of medium heights. At the same time, Wild jack (*A. hirsutus*) bears multiple oval/ellipsoidal, fleshy, smaller, orange/yellow sheathed fruits of size 6-10 cm at the apices of branches of higher heights. Such diverse phenotypic characters suggest differentiated pollinator and disperser networks and differential mechanisms (Jarrett, 1977; Matthew et al., 2006; Jagtap and Bapat, 2010; Ragone, 2018; Buddhisuharto et al., 2021).

**Figure 1:**
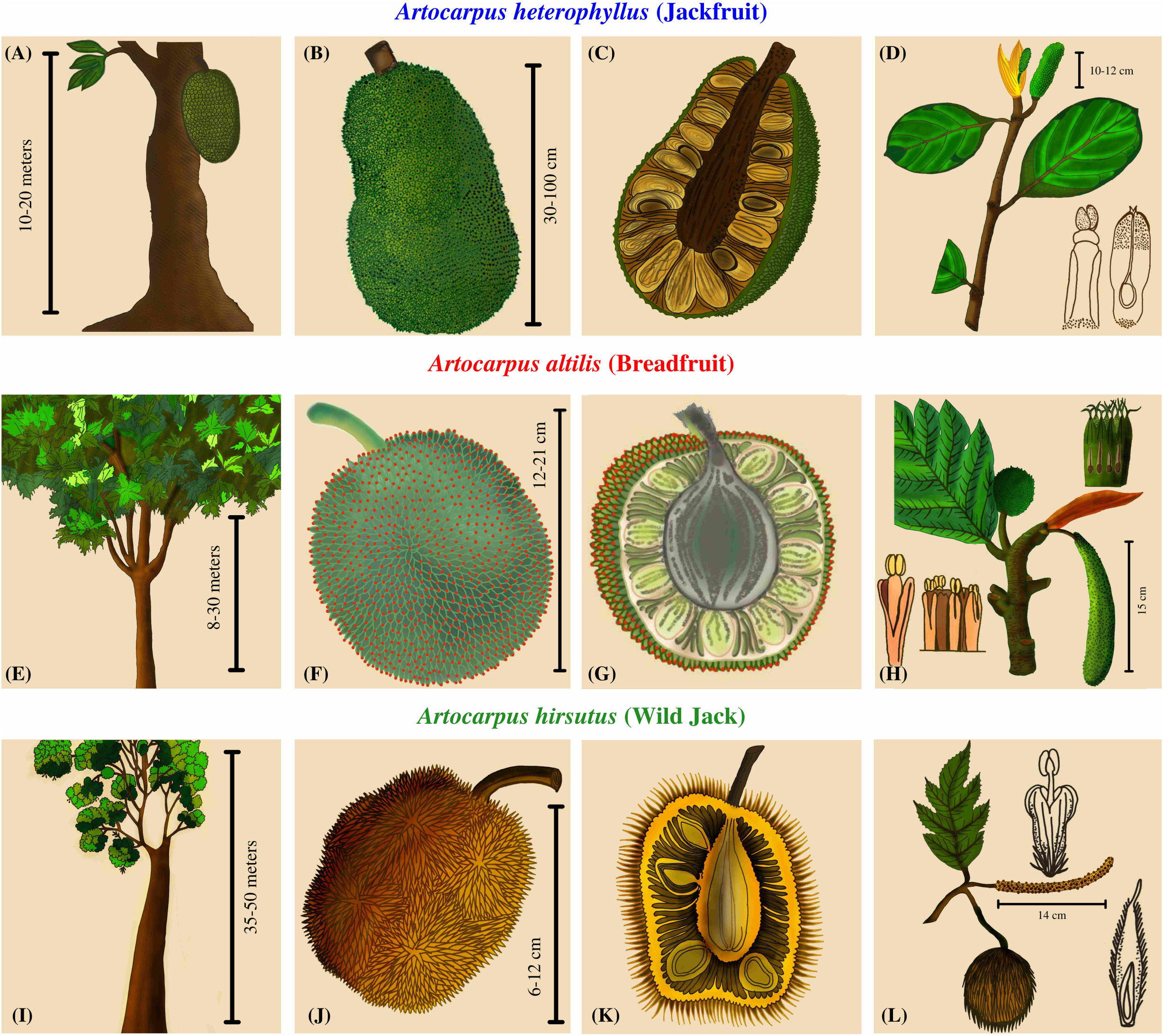
Plant morphology and comparative phenotypic characters of Artocarpus species. **A, E, and I**: The first column depicts the overall structure of the trees. **B, F, and J**: The second column contains drawings of the fruit describing the color, shape, and size. **C, G, and K**: The third column has a cross-sectional view of the fruits depicting the number and arrangement of seeds. **D, H, and L**: The fourth column has drawings of the inflorescences.

The Wild Jack (*A. hirsutus*) is unique in its phenotype compared to the other two popular Artocarpus species. Due to its endemic distribution in the Western Ghats and its forests, it has received limited attention and is still understudied (Matthew et al., 2006). However, in its native range, it’s a multipurpose plant of economic and ecological importance. Its long, high-quality, pathogen-resistant timber is widely used for building houses, boats, and large, long-lasting structures (Matthew et al., 2006; Xavier et al., 2014; Meenu et al., 2021). Its other parts, like fruits and seeds, are used as a rich source of energy and taste. Leaves, seeds, and bark are used in traditional medicinal treatments (Matthew et al., 2006; Jagtap and Bapat, 2010; Solanki et al., 2020; Buddhisuharto et al., 2021; Meenu et al., 2021). The constituents of this fruit are comparable or superior to the other two species, which demands further research into this plant (Solanki et al., 2020). Wild jack reaches maturity for timber harvesting in around 20 years, and the present distribution of the species cannot fulfil the increased demand of the timber market, which makes it vulnerable to population decline. Hence the re-assessment of conservation status and efforts to effectively conserve this plant is warranted (Matthew et al., 2006). The sacred groves of the Western Ghats are fragmented forests protected by locals due to religious importance. Wild jack is a ubiquitous constituent of these ancient forests providing some conservation for this species (Chandrashekara and Sankar, 1998; Tambat et al., 2005). But recently, these forests have been threatened due to deforestation and developmental projects (Matthew et al., 2006; Osuri et al., 2014). The biodiversity hotspot of the Western Ghats is home to multiple endemic flora and fauna (Myers et al., 2000), but there is a paucity of genomic data from these species. By generating genomic resources, our study aims to incorporate Wild jack in a comparative framework with other Artocarpus species. Hence, our goal is to identify genomic changes related to differential phenotypes and acclimatization to their habitats. In line with this goal we envisage the following objectives for this study:

1. Sequence and assemble the Wild jack genome (and plastome), i.e., *Artocarpus hirsutus*.
2. Perform gene annotation to identify orthologous gene sequences across order Rosales members and construct a species tree.
3. Analyse gene family evolution in these species, especially in Artocarpus members.
4. Use demographic reconstructions to analyse population size history and species distribution modelling to assess species range dynamics for Artocarpus species.
5. Detailed repeat sequence annotation to understand repeat accumulation dynamics.
6. Employ multiple methods to detect the signatures of selection in the genes of all Artocarpus species.

## Materials and Methods

### Sample Collection, genome sequencing, and assembly

*Artocarpus hirsutus* is endemic to the Western Ghats and its forests. We located a fruit-bearing tree near the College of Forestry, Ponnampet (GPS coordinates 12°08′56.5″N 75°54′32.5″E; Altitude: 829–850 m Above Sea level) and sampled some leaves for the sequencing. A leaf was cleaned, sanitised, and then cut into pieces for further processing. The whole genomic DNA was extracted from the leaves using QIAGEN’s DNeasy plant mini kit. The quality of the extracted DNA was evaluated by observing the DNA band on 1% Agarose gel for shearing. The concentration and purity of extracted DNA were assessed using QUBIT 3.0 and Nanodrop. The purified DNA was then used to prepare Illumina short-read (150 bp) libraries with TruSeq DNA Nano Library Prep Kit with an insert size of 450 ± 50 bp. We sequenced ~ 88X coverage of genome with Illumina short-read paired-end data using the Illumina Novaseq 6000 sequencer.

The quality of whole-genome sequencing (WGS) paired-end reads was assessed using FASTQC. Barcode sequences were trimmed if present using Cutadapt (Martin, 2011). These sequencing reads were used for the estimation of genome size. The genome assemblies of previously published congeneric species vary from ~800 Mbp (*A. altilis*) to ~980 Mbp (*A. heterophyllus*); their Kmer-based genome size estimates are 812 Mbp and 1005 Mbp, respectively (Sahu et al., 2019). We used Jellyfish (Marçais and Kingsford, 2011) to perform Kmer analysis using kmer-size (k) of 21 and hash size (-m) of 100M on the sequencing reads.

GenomeScope (Vurture et al., 2017) was then used to estimate genome size and heterozygosity. The 21 kmer value-based genome size estimate was 635.16 Mbp with 1.16% heterozygosity. These estimates show that *A. hirsutus* has a slightly smaller genome size than other congenerics. We used the Celera assembler implemented in MaSuRCA version 4.0.6 (Maryland Super Read Cabog Assembler) for assembling the sequencing data (Zimin et al., 2013). The published assemblies of *A. heterophyllus* and *A. altilis* were used as a reference for the synteny-assisted assembly step of MaSuRCA. The resultant genome assembly was 791.16 Mbp in length. We used Quast (Gurevich et al., 2013) to calculate genome assembly metrics such as N50 and L50 (**Supplementary Table 1**). BUSCO version 3 (Simão et al., 2015) was used to assess the genome completeness with the eudicotyledons_odb10 dataset (**Supplementary Table 2**).

### Repeat annotation and analyses

For de-novo identification and annotation of the repetitive genomic regions and/or transposable elements, we used RepeatModeler version 2 (Flynn et al., 2020). We used the LTR_struct option to include LTR models identified by programs such as LTR-FINDER (Xu and Wang, 2007) and LTR-Harvest (Gremme et al., 2013). The consensus fasta library obtained by RepeatModeler 2 was then used as input to RepeatMasker (Smit, AFA, Hubley, R & Green) to annotate, mask and tabulate the repeat content and their types. The resultant output file was then used to soft mask the genome for further analyses. The RepeatMasker .align output was used to calculate Kimura two-parameter divergence estimates (TE age) between the repeat families for all species using accessory scripts provided with RepeatMasker suite like buildSummary.pl, calcDivergenceFromAlign.pl, and createRepeatLandscape.pl. The obtained output was summarised to plot histograms of Kimura divergence values to visualise the distribution of repeat families across the time scale.

### Genome annotation

We used MAKER version 2 (Campbell et al., 2014) to annotate the genome’s coding regions. Three rounds of the maker pipeline were executed to obtain the final annotated genesets. In the first round of homology-based annotation, we used protein fasta sequences of species of order Rosales (*Artocarpus altilis, Artocarpus heterophyllus, Morus notabilis, Parasponia andersonii, Trema orientale, Ziziphus jujuba, Rhamnella rubrinervis, Cannabis sativa*) from NCBI (**Supplementary Table 3**). The mRNA evidence from *A. altilis* was provided as alternative transcript sequences. The obtained genesets from this round were then used to generate training gene models for de-novo gene annotation algorithms like SNAP (Johnson et al., 2008) and AUGUSTUS (Stanke and Morgenstern, 2005). New gene models were identified during both rounds, and existing gene models were refined. Genesets after the third round were considered final and used to get coding sequences and translated protein sequences.

### Chloroplast assembly, annotation, and analysis

The chloroplast sequence was independently assembled using WGS reads with NOVOPlasty version 4.3.1 (Dierckxsens et al., 2017). The chloroplast genome sequence of *A. altilis* (NCBI accession: NC_059002.1) was used as a reference for the algorithm, and the Maturase K gene sequence of *A. hirsutus* (NCBI accession: KU856362.1) was used as a seed sequence, which is used as sequence generation point for the assembly. The resultant assembly produced two contigs with only one arrangement possibility leading to a complete circular genome sequence spanning ~162Kbp. The chloroplast assembly was then annotated using GeSeq (Tillich et al., 2017), and the circular genome was depicted and visualised using OGDRAW (Greiner et al., 2019) implemented in CHLOROBOX. Currently available chloroplast genomes from the Artocarpus genus and outgroup species *Ficus religiosa* and *Morus indica* were downloaded from NCBI. To investigate rearrangements between these chloroplast genomes, they were aligned with ProgressiveMauve aligner (Darling et al., 2010) and visualised in Mauve alignment viewer (Darling et al., 2004). To identify the phylogenetic positions of these genomes, we aligned the genomes using the MAFFT aligner (Katoh et al., 2002). The appropriate substitution model was estimated using Modeltest-ng (Darriba et al., 2020), and the phylogenetic tree was constructed using Raxml-ng (Kozlov et al., 2019) with 1000 bootstraps. The chloroplast genomes of *A. heterophyllus* and *A. integer* show an inversion for the SSC (Small Single Copy) region compared to other Artocarpus sp. Plastomes (**Supplementary Figure 1**).

### Identification of orthologous sequences and construction of species tree

The translated protein sequences of *A. hirsutus* and 13 other species (*A. altilis, A. heterophyllus, M. notabilis, P. andersonii, T. orientale, C. sativa, R. rubrinervis, Z. jujuba, M. baccata, M. domestica, P. persica, F. vesca, and R. chinensis*) were concatenated and used to find orthologs. We used Orthofinder (Emms and Kelly, 2019) to find orthologous genic sequences across 14 species with parameters to use MSA alignments to obtain the orthogroups using diamond blast (Buchfink et al., 2021), MAFFT (Katoh et al., 2002) and fasttree (Price et al., 2009). The orthologous gene sequences in which *A. hirsutus* is present were tabulated. These gene IDs were used to get corresponding CDS sequences from each species. These CDS sequences for all the genes were then aligned using GUIDANCE version 2 (Sela et al., 2015) with the MAFFT aligner (Katoh et al., 2002). All the resultant CDS alignments were concatenated and used to find partitions and models using IQTREE version 2 (Minh et al., 2020). After that, the loci and concatenated trees were obtained to get bootstrap support with additional metrics such as Gene concordance factor (gCF) and Site concordance factor (sCF). Following these evaluations, the tree was exported and was rooted at a branch leading to *F. vesca* and *R. chinensis*.

### Comparative genomics and gene family analyses

The proteomes of 4 species, *A. hirsutus, A. altilis, A. heterophyllus*, and *M. notabilis* were used to identify overlapping and non-overlapping gene clusters using Orthovenn version 2 (Xu et al., 2019). Orthovenn identified and constructed the unique and common gene clusters for all four species. Unique gene clusters of Artocarpus species were subjected to GO enrichment analysis. We used CAFÉ version 5 software (Mendes et al., 2021) for gene family analyses of contractions and expansions. We first concatenated proteomes of 14 species used for species tree construction and made a blast database. This 14-species proteome database was used as a subject to perform all vs. all protein blast (blastp) (Camacho et al., 2009). The blast results were then used as input for mcxload to create network and sequence dictionary files. The clustering was performed using mcl clustering software (Li et al., 2003) with an inflation parameter (-I) of 3. The cluster files were then reformatted, and the gene families with large gene copy numbers were removed from the analyses. The constructed species tree was converted to an ultrametric tree using r8s software (Sanderson, 2003) using a divergence estimate of 87 MYA between *P. persica* and *Z. jujube* obtained from TimeTree (Kumar et al., 2017). The filtered clustering file of MCL and the ultrametric tree were then used as input for the CAFÉ 5. The clade-based gene family expansion/contraction results were then summarised and represented on the phylogeny. Out of all significant gene family contractions, we selected only those gene families with a difference of five gene copies at the least between the species. By enforcing these stringent criteria, we got seventeen, three, and seven gene families significantly expanded in *A. hirsutus, A. heterophyllus*, and *A. altilis*, respectively.

### Lineage-specific selection tests in Artocarpus genes

Tests of selection intensity among species for the same orthologous genes in a phylogenetic framework provide opportunities to identify loci under relaxed or intensified pressures in a focal species of interest. This selection pressure analysis helps us identify evolutionary changes and signs of adaptations to their bioclimatic niche. We used multiple approaches to identify selection pressures in Artocarpus to understand the evolutionary mechanisms and processes this plant has undergone. We used branch-site models implemented in PAML version 4.9 (Yang, 2007) and aBSREL (Adaptive Branch-site Random Effects Likelihood) (Smith et al., 2015) implemented in HYPHY to test for positively selected branches. We also used RELAX (Wertheim et al., 2015) (intensification parameter; K > 1) implemented in HYPHY to identify the genes under intensified selection. For detecting strong purifying or relaxed selection, we implemented the branch site model of PAML version 4.9 and RELAX (relaxed parameter; K < 1) of HYPHY. To reduce the false positive results, we compared the list of genes identified as positively selected by all three methods and considered only those genes that were overlapping/common between them. The functional roles of positively selected genes were cross-referenced using KEGG (Kanehisa and Goto, 2000) and FLOR-ID (Bouché et al., 2016) databases.

### Demographic history reconstruction

The genomic sequencing reads of one individual each of *A. hirsutus, A. altilis*, and *A. heterophyllus* were mapped to the respective genome assemblies using the BWA MEM aligner (Li, 2013). The alignments were converted to binary, sorted, and indexed using samtools (Li et al., 2009). These binary alignments were then used to call consensus sequence using bcftools (Li and Barrett, 2011). To assess the effect of genomic regions such as exonic, intronic, intergenic, and repetitive elements on the demographic estimation, we masked each part to check the impact of the respective fraction. We masked the respective genomic region using BEDTOOLS maskfasta and followed similar steps mentioned above to get the demographic estimation. To assess the effect of each individual repeat family/type, we followed the published protocol/scripts (Patil and Vijay, 2021). We used filters like -C50, - Q30, -q 20 for bcftools mpileup to ensure quality bases and mapped reads to be considered in the variant calling. The consensus calls were converted to the required (fastq) format using vcfutils vcf2fq using a quality filter of 25, whereas calls with less than one-third and more than twice the mean coverage were excluded during this step to exclude false calls. These consensus calls were then converted to input format (.psmcfa) for psmc using fq2psmcfa. The input psmcfa file was then used to run the psmc program (Li and Durbin, 2011) with options - N 25 -t 5 -r 5 -p 4+25*2+4+6. The output of psmc was inspected for a sufficient number of recombination events. At first, we used a mutation rate (μ) of *Populus trichocarpa*, i.e., 2.5e-09 per site per year (Tuskan et al., 2006) with a generation time of 15 years to execute the psmc_plot.pl script to get scaled demographic trajectories for each species.

### Estimation of mutation rate

Since we were trying to study demographic effects on these three species comparatively, we needed to understand the bottleneck events for these species from their native ranges. *A. hirsutus* and *A. heterophyllus* are native to the Western Ghats and should have experienced similar demographic events. Our initial scaled PSMC plots for both species with the same mutation rates did not align with the starting point of the trajectory. These trajectories created a possibility that there might be mutation rate differences between these three species. To obtain a reliable mutation rate estimate, we sampled orthologous alignments in which only four species (*A. hirsutus, A. altilis, A. heterophyllus*, and *M. notabilis*) are present. We fixed an input un-rooted tree structure to allow branch-specific comparisons possible. We used codon alignments of ~1500 genes to estimate a (d_4_) 4-fold degenerate site substitution rate (parameters used, model = 0, NSsites=0, seqtype=1, CodonFreq=2, runmode=0) using PAML version 4.9. The obtained d_4_ rates for all alignments were summarised, and the mean value for these estimates was considered d_4_ for individual species. These mean d4 estimates were then divided by the divergence time between compared branches or species. The estimates obtained were then considered a proxy of the respective species’ mutation rates (Nadachowska-Brzyska et al., 2015).

### Species Distribution Modelling

We downloaded species occurrence data corresponding to the native range of each species as identified earlier (Williams et al., 2017) from the GBIF (Global Biodiversity Information Facility) database for all three Artocarpus species (Gbif.Org, 2022). We used the method of Ecological niche modelling (ENM) to predict the species distribution during three paleoclimatic eras: Last Glacial Maximum (LGM, approx. 20,000 years ago), Last interglacial (LIG, approx. 110,000-130,000 years ago), and Marine Isotope Stage 19 (MIS19, approx. 750,000-790,000). The environmental variables for these periods were extracted from the database PaleoClim (Brown et al., 2018) at a resolution of 2.5 min arc. Environmental layers were resized to the species’ native range using the software DIVA-GIS (version 7.5) (Hijmans et al., 2012). We considered the annual estimates and excluded the seasonal variables for highly correlated bioclimatic variables. The set of variables used was chosen based on species-specific considerations for the compared periods (see **Supplementary Table 4B-C**).

The ENM was performed using the software MaxEnt (version 3.4.4). The MaxEnt software has a wide variety of modelling settings and is particularly popular due to its high predictive performance in species distribution modelling. The model minimises the relative entropy between two probability densities; one probability estimated from the presence-only data and one from the landscape (Elith et al., 2011; Merow et al., 2013). The settings for MaxEnt were species and paleoclimatic era-specific. We used the R package ENMeval, which identifies settings that balances model fit and increases the predictive ability (see **Supplementary Table 4E**) (Muscarella et al., 2014). The following settings were set by default: 10000 background points, 500 maximum iterations, ten runs of cross-validations, and the regularisation multiplier were explicitly based on ENMeval results. We saved the output in cloglog form, which is the simplest to understand and the default output format. It gives the probability of occurrence estimate between 0 to 1. We selected the mean of all ten replicate runs to represent each species across each time period. The accuracy in the prediction of species distribution was analysed through the use of a receiver operating characteristics (ROC) plot. In the ROC plots, all the values fell between 0 and 1 (AUC). All the values were above 0.5 and are considered better than random when the curve lies above the diagonal, indicated by the area under the curve (AUC) (see **Supplementary Table 4A**) (Merow et al., 2013). A Jackknife test was performed to find the different contributions of variables and to identify the ones with a maximum contribution (see **Supplementary Table 4D**). The habitat suitability maps of species distribution were generated using R.

## Results

### Lineage-specific gene family dynamics

Gene family expansion and diversification are prominent drivers of phenotypic evolution. Comparative analysis of Artocarpus genomes and outgroups from Rosales’ order identified changes in gene family composition (see **Figure 2A, B** and **Supplementary Table 5, 6**). Notably, lineage-specific genes identified by orthovenn2 in *A. hirsutus* are enriched for pollen recognition genes (GO: 0048544), which are Receptor Kinases from the Lectin domain-containing gene family (**Supplementary Table 6**). We also found evidence of lectin gene family expansion based on CAFÉ analysis in *A. hirsutus* (41 copies) compared to *A. altilis* (33 copies) and *A. heterophyllus* (22 copies) (see **Figure 1C**, RK3 panel, **Supplementary Table 7**). Among the lectin genes with orthologs across all three Artocarpus species, we detect signatures of intensified selection using RELAX and positive selection using PAML branch-site and aBSREL (**Supplementary Figure 2**). Lectin domain-containing proteins have diverse functions in biotic and abiotic stress response, plant growth, and development (Sun et al., 2020; Saidou and Zhang, 2022). Our results suggest diversification of lectin domain-containing proteins in *A. hirsutus*.

**Figure 2:**
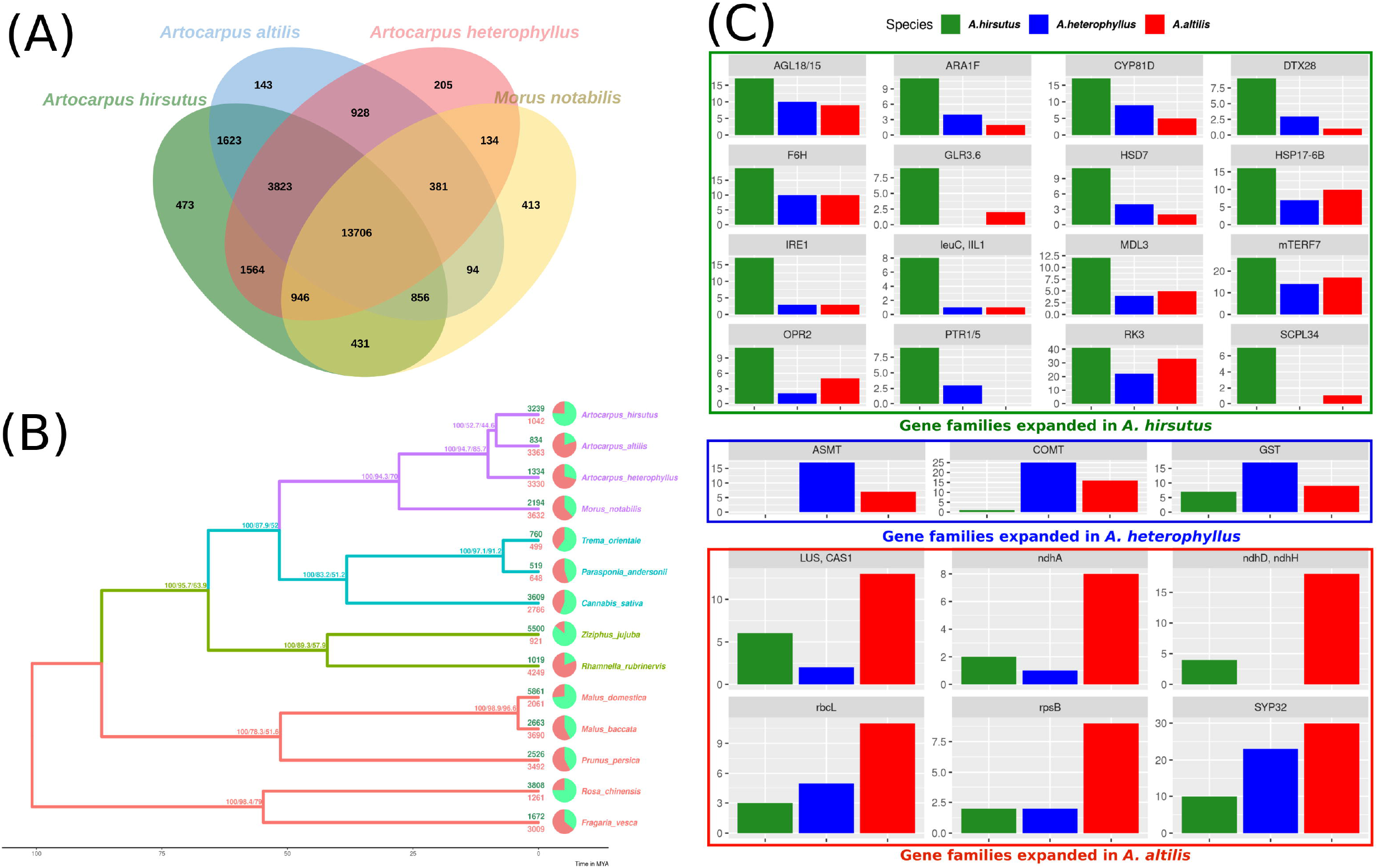
Orthologous relationship between Artocarpus and their relatives. **A.** Orthovenn diagram of three Artocarpus species (*A. hirsutus, A. altilis*, and *A. heterophyllus*) with *M. notabilis*. Unique and shared gene clusters between these species are denoted in respective intersections. *A. hirsutus* has 473 unique gene clusters not shared with other congenerics. **B.** Representative species tree with gene family evolution or Gene gain-losses: 14 species from Order Rosales of family Moraceae, Rhamnaceae, Cannabaceae, and Rosaceae are represented in the species tree based on ~4500 orthologous genes. Branch values are bootstraps values, gCF (gene concordance factors and sCF (site concordance factors), respectively. Pie charts and associated numbers at the tips show gene families’ expansions (green) and contractions (red). **C.** Significantly expanded gene families in *A. hirsutus* (green), *A. heterophyllus* (blue), and *A. altilis* (red).

Apart from lectins, *A. hirsutus* showed lineage-specific gene family expansions in at least 15 other gene families with functions varying from pollen/flower development (AGL18/15, ARA1F, PTR3, EDA17), secondary metabolite biosynthesis (F6’H, IIL1, HSD7, DTX28 (Upadhyay et al., 2020), MDL3), stress tolerance and defence, i.e., biotic (IRE1, IIL1, DTX28, MDL3) and abiotic (F6’H, CYP81D8, GLR3.6, SCPL34, PTR1/5, HSP17-6B, OPR2), growth and development (mTERF7, HSD, AGL18/15, ARA1F) and plant-pathogen interactions (IRE1, F6’H, IIL1, OPR2, MDL3). All these numerous gene family expansions may reflect the concerted evolution of this plant to acclimatise to biotic and abiotic conditions and adapt to its habitat.

In *A. heterophyllus*, the lineage-specific genes identified by orthovenn2 are enriched for Toll-Interleukin-Resistance (TIR) domain proteins, Receptor Like Protein 33 (RLP33), and the flavonoid biosynthesis pathway. TIR domain proteins and RLP33 are well known for foreign pattern recognition and providing immunity to plants from microbes (Burch-Smith and Dinesh-Kumar, 2007; Jamieson et al., 2018). In addition, the two most essential genes of the Flavanoid Biosynthesis Pathway, Chalcone Synthase (CHS) and Flavanone 3-Hydroxylase (F3H) have lineage-specific gene copies and may explain the high flavonoid content of *A. heterophyllus* (Meera et al., 2018). ASMT (N-Acetylserotonin Methyltransferase) and COMT (Caffeic Acid O-methyltransferase) genes act in the penultimate step of the melatonin pathway (Back et al., 2016; Zhao et al., 2019). COMT also plays an important role in the lignin biosynthesis pathway (Wang et al., 2013). The copy number of both ASMT and COMT genes is higher in *A. heterophyllus* (17 and 25 copies) compared to *A. hirsutus* (0 and 1 copies) and *A. altilis* (7 and 16 copies) (**Supplementary Figure 3**). The gene family of Glutathione S-Transferases (GST), which have a role in stress tolerance, has also expanded in *A. heterophyllus*.

The lineage-specific genes of *A. altilis* are enriched for Hexokinase-3, ABCB27 (ATP-Binding Cassette B27) or ALS1 (Aluminium Sensitive 1), and mTERF15. Hexokinase-3 is involved in sugar processing, primarily glucose and development (Paulina Aguilera-Alvarado and Sanchez-Nieto, 2017). ABCB27 or ALS1 are transporters involved in stress response to Aluminium-rich or Acidic soils (Kar et al., 2021). The mTERF15 is a transcription factor which regulates the expression of mitochondrial assembly factor I genes, specifically NAD2/3 (NADH ubiquinone oxidoreductases), and regulates energy generation (Hsu et al., 2014). Interestingly, we found gene family expansions in multiple genes of mitochondria and chloroplast (ndhH, ndhD, ndhA, rbcL, and rpsB) in *A. altilis*. These expansions of organellar genes and their regulators might be due to a higher energy requirement caused by oxidative stress or other stressors. The triterpenoid biosynthesis synthase genes like Cycloartenol Synthase (CAS1) and Lupeol synthase 2/5 (LUP2/5) (Thimmappa et al., 2014; Cárdenas et al., 2019) and essential pollen development proteins, syntaxin of plants (SYP31/32) (Rui et al., 2021) were also increased in copy number.

### Demography rather than phylogeny determines the population histories

We found that *A. altilis* underwent demographic contraction ~ 2 to 1 million years ago (MYA), followed by extensive population expansion from ~ 1 MYA to 150 thousand years ago (KYA), which marks the start of the Holocene or Last Glacial Period (**Figure 3C**, and **4A**). In contrast to *A. altilis*, the population sizes of *A. heterophyllus* and *A. hirsutus* experienced extensive expansion from ~ 2 to 1 MYA, followed by a contraction in population size from ~ 1 MYA to 200 KYA (**Figure 3A-B**, and **4A**). After the onset of the Holocene, the effective population size declined in *A. altilis* and *A. heterophyllus*. However, in *A. hirsutus* the population size recovered and stabilised before undergoing another round of population decline in the Holocene. Comparing demographic histories among the three Artocarpus species and the species distribution models suggests that bioclimatic changes and the habitat have been instrumental in shaping the population histories. In conclusion, the demographic histories of the Artocarpus species reflect the effects of habitat more than their phylogenetic relatedness (**Figure 4A**).

**Figure 3:**
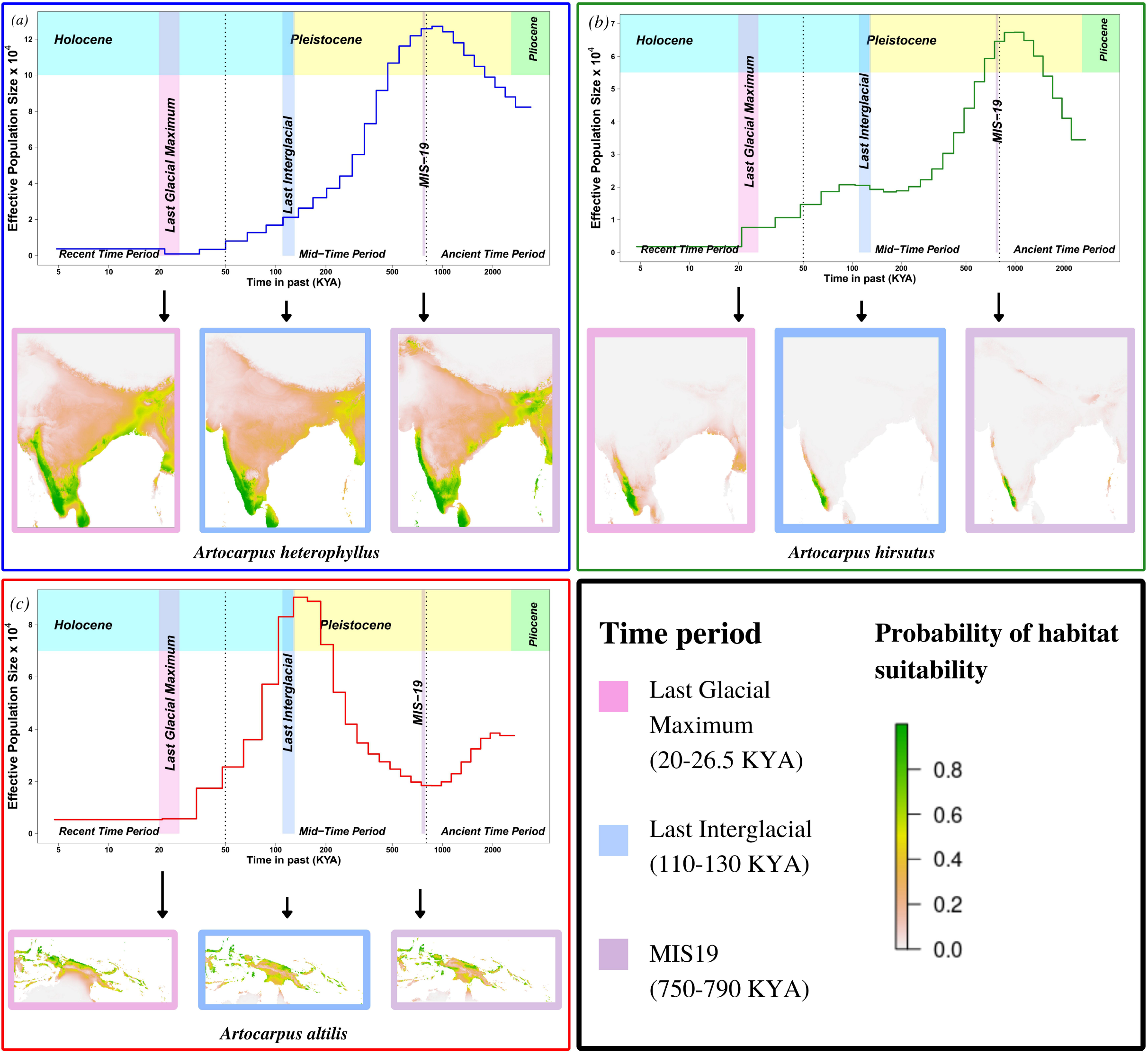
Demographic history reconstruction and species distribution modelling of Artocarpus species. Demographic history reconstruction using PSMC with Species distribution modelling for LGM, LIG, and MIS19 for **A.** *A. heterophyllus* (blue), **B.** *A. hirsutus* (green), **C.** *A. altilis* (red).

**Figure 4:**
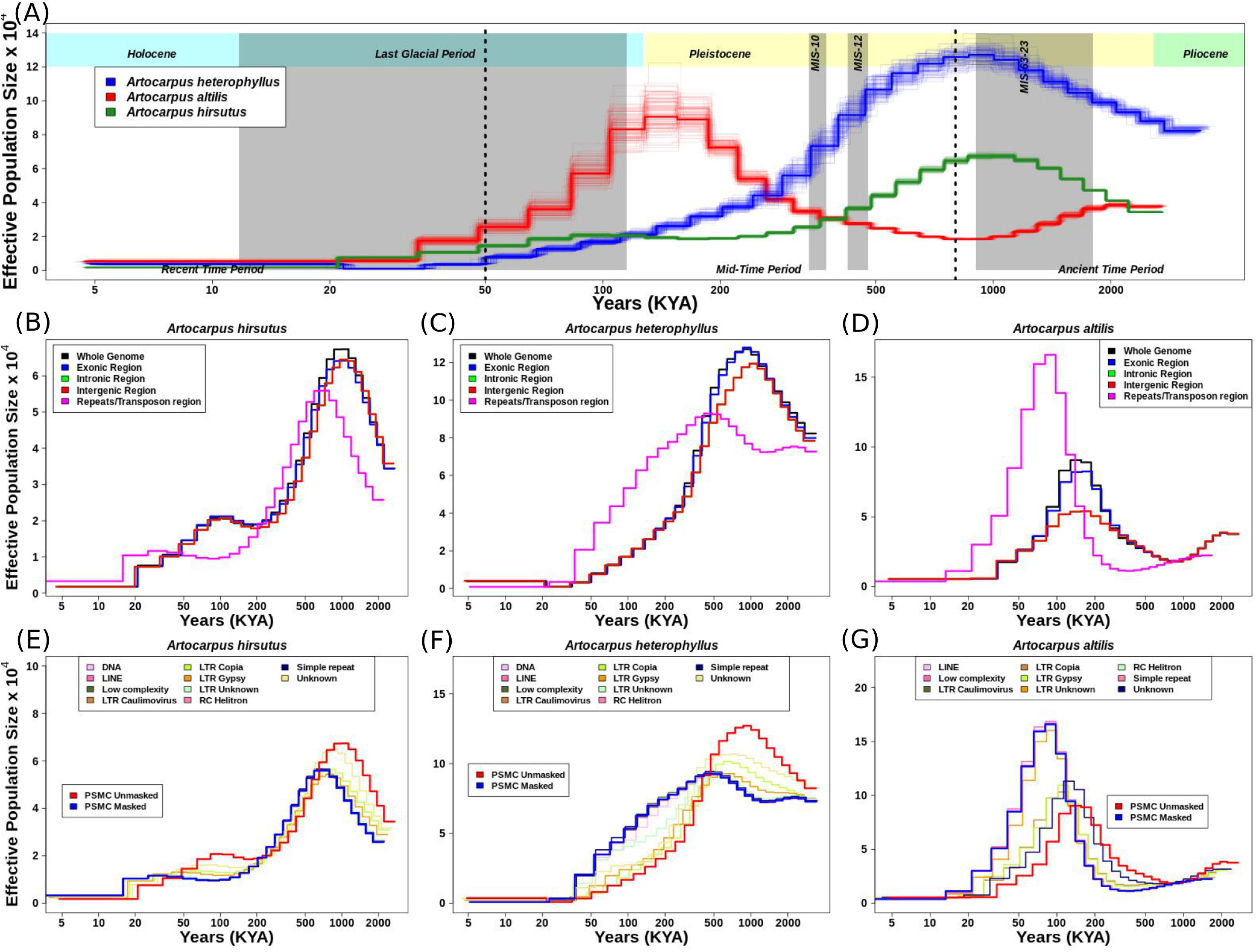
Demographic history reconstruction of three Artocarpus species. **A.** Demographic history of *A. heterophyllus* (blue), *A. altilis* (red), and *A. hirsutus* (green) are represented by the respective trajectories with additional bootstrapped lines. *A. heterophyllus* and *A. hirsutus* show similar trends in population size changes during the Pleistocene, which might be due to similar distribution in the Western Ghats and hence, similar demographic processes for both species. Even though *A. altilis* is phylogenetically closer to *A. hirsutus;* the demographic trajectory is drastically different compared to the other two Artocarpus species, indicating the effects of difference in its distribution. Demographic trajectories and the effect of genome fraction used for PSMC inference in **B.** *A. hirsutus*, **C.** *A. heterophyllus*, and **D.** *A. altilis*. The genomic fractions used are the Exonic region, Non-exonic (Introns and Intergenic) region, Repetitive genomic regions, and whole genomic region. In all three species, repeat regions show the most drastic changes in the inferences. The effect of different repeat types on PSMC inference is represented in **E.** *A. hirsutus*, **F.** *A. heterophyllus*, and **G.** *A. altilis*. All three species show differences in trajectories at various levels. But LTR classes like LTR-Copia, LTR-Gypsy, and unknown repeats seem to drive most of the changes in the trajectories in all species. The amount of change due to LTR-Copia is most prominent in *A. heterophyllus* regarding the trajectory. However, *A. altilis* experienced the most noticeable change in the trajectories across the x-axis, which can be a consequence of differences in mutation rate.

The estimates of historical effective population size (N_e_) reflect evolutionary processes such as actual changes in population size, population structure, gene flow (Mazet et al., 2015, 2016), and linked selection (Schrider et al., 2016) and/or regions of the genome used (Patil and Vijay, 2021). Hence, we evaluated the effect of using different genomic regions in estimating demographic histories. Exon, intron, and intergenic region-masked trajectories matched with the whole-genome-based trajectory, which explains that these individual regions of the genome are not drastically changing the estimates (**Figure 4B-D**). However, masking repetitive genomic regions resulted in two types of changes in the inferred trajectory. The less noticeable trajectory change results in a diagonal shift towards recent time intervals in *A. hirsutus* and *A. altilis*. The most drastic change in trajectory occurs in *A. heterophyllus*, where the repeat masked and whole-genome-based N_e_ estimates lack concordance. Our results indicate that repetitive genomic regions are the most notorious factor affecting demographic estimation in Artocarpus species.

To understand which type of repeat regions affect the inference of demographic history in these species, we investigated the effect of each repeat type. In all three species, the shift in trajectories among LTRs (i.e., LTR-Unknown, LTR-Gypsy, and LTR-Copia) was highest (**Figure 4E-G**). Other repeat families, like simple repeats, DNA transposons, low complexity regions, etc., mirrored the masked trajectory and had no effect of masking on demography. LTR-Gypsy repeat family had the most noticeable impact on the trajectories of *A. hirsutus* and *A. altilis*, whereas LTR-Copia greatly impacted the N_e_ estimates of *A. heterophyllus*. The most surprising result of repeat masking occurs in the N_e_ estimates of *A. heterophyllus*, which drastically changes the trajectory in magnitude and shape mainly due to LTR-Copia.

### Differential abundance/accumulation of repeat families in *A. heterophyllus*

The repetitive genomic regions strongly affected the demographic analyses, which demands further detailed characterisation of repeat families and their contents. We compared the types of repeat families assembled in the Rosales genomes and their abundances (see **Figure 5A**). Of the three Artocarpus genomes, *A. hirsutus* (481 Mbp; 60.51% of the genome) and *A. altilis* (505 Mbp; 60.68% of the genome) have a comparable composition of repeat types. In contrast, *A. heterophyllus* (614 Mbp; 62.56% of the genome) has a higher overall repeat content than the other two species. Specifically, the abundance of LTR-Copia in the *A. heterophyllus* genome (246.5 Mbp; 25.1% of the genome) was highly elevated compared to *A. altilis* (131 Mbp; 15.7% of the genome) and *A. hirsutus* (128 Mbp; 16.1% of the genome). Other than LTR-Copia, most other families, except for some unknown/unannotated LTRs, were similarly abundant across the three Artocarpus species. These differences in repeat composition suggest a species-specific expansion or excessive accumulation of LTR-Copia family repeats in *A. heterophyllus*. Cannabis sativa has LTR-Copia and overall repeat content expansion similar to *A. heterophyllus* among the Artocarpus outgroup genomes.

**Figure 5:**
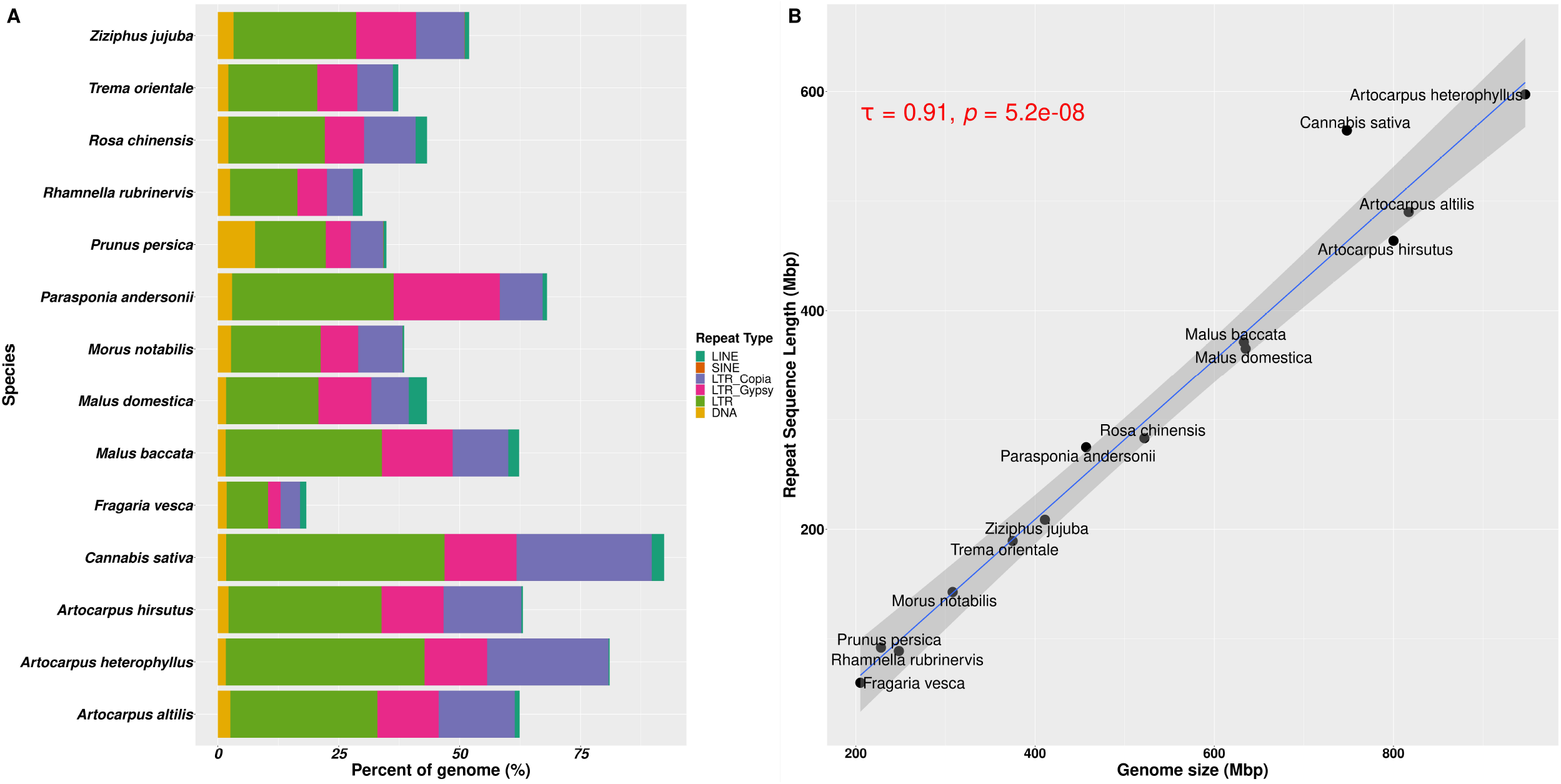
Repetitive genomic regions and their abundance in Rosales genomes. **A.** Repeat content percentages in genomes of 14 species of Rosales with the abundance of types of repeats across these species. **B.** Correlation of genome sizes with total repeat sequences assembled in these 14 genomes. There is a strong positive correlation (τ = 0.91 and p-value = 5.2e-08) between the abundance of repeat sequences assembled in the genome and the assembly sizes of these 14 species.

Genome size evolution is a product of various factors, including the repetitive profile of the species. Repeat sequence accumulation can inflate the genome size of a species and shape genome evolution. To address if repeat expansions and assemblages in the genomes of Order Rosales significantly impacted their genome sizes, we correlated their total assembly sizes (i.e., a proxy for genome size) and assembled repeat content. We observed a strong positive correlation between total assembled repeat length in these genomes with their genome sizes (**Figure 5B**, Kendall’s correlation coefficient = 0.91, p-value = 5.2e-08). The strong correlation suggests that Order Rosales underwent genome size evolution strongly influenced by repeat expansions and accumulations.

To understand the differential species-specific repeat accumulation in the Rosales order, we used Kimura two-parameter divergence estimates to reconstruct the timeline of repeat expansion (see **Figure 6**). All three Artocarpus species have a comparable repeat abundance of ~2% genomic content at a Kimura distance of ~0.1 and likely represent their shared history of repeat accumulation. However, *A. heterophyllus* has recently accumulated species-specific repeats corresponding to ~3.5% genomic content, primarily LTR-Copia sequences at a Kimura distance of ~0.05. Hence, the recent accumulation of the LTR-Copia is most likely the reason for genome size expansion in *A. heterophyllus* after divergence from *A. altilis* and *A. hirsutus*. Like *A. heterophyllus, Cannabis sativa* also has a similar pattern of recent LTR-Copia repeat accumulation compared to other members of Cannabaceae. Rosaceae family has a rich diversity of plants with flowers and fruits with economic and commercial value. Plants of this family prove to be diverse in terms of species-specific repeat type accumulation. For instance, Malus species have a recent expansion of LINE sequences.

**Figure 6:**
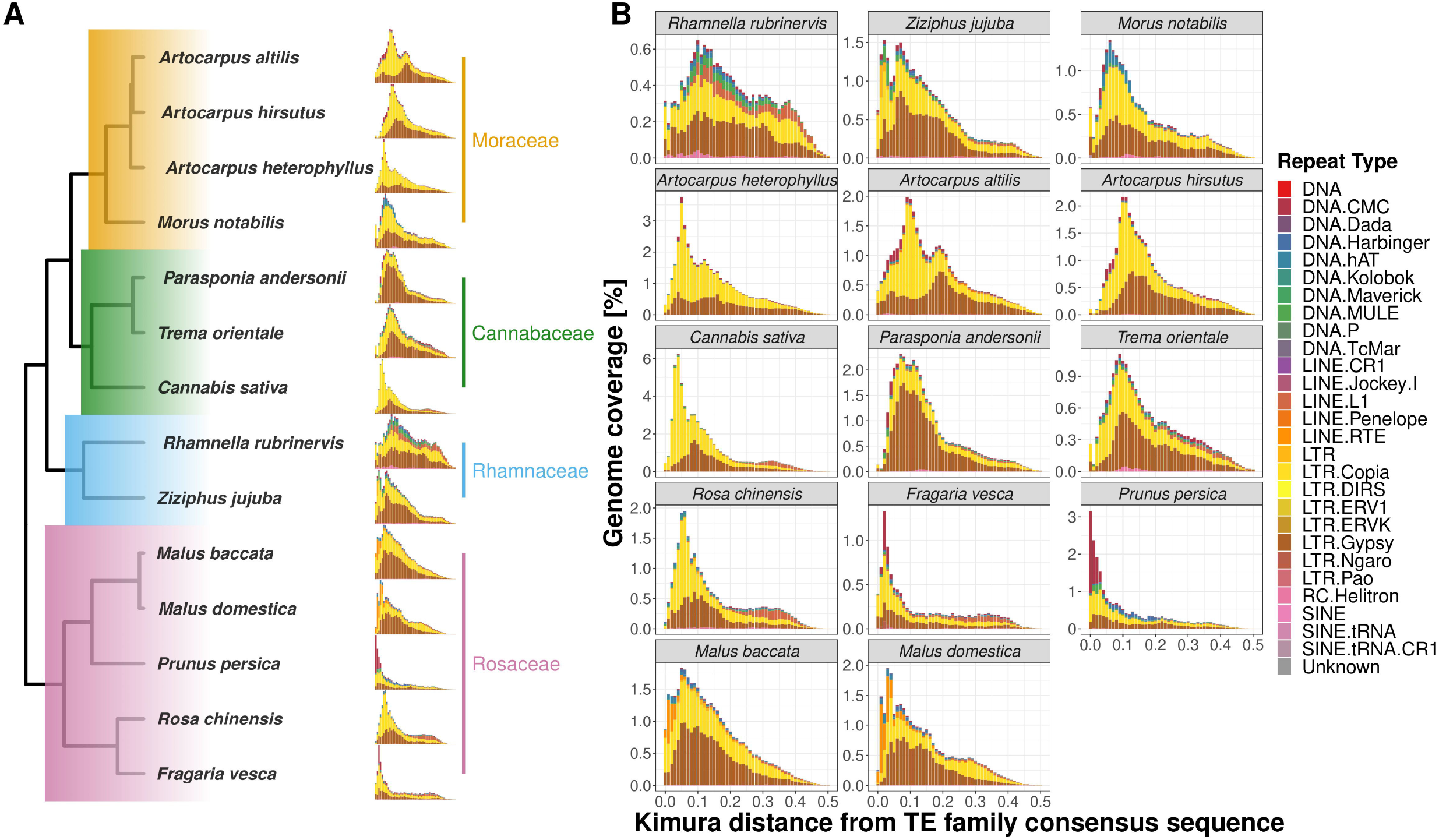
Repeat landscape evolution in 14 species of order Rosales. Family-wise changes in repeat landscapes: **A.** The differences in abundance and insertion ages of repeat families are depicted phylogenetically. The recent spike in the fraction of the genome covered by repeats of *A. heterophyllus* suggests a recent accumulation not shared by the other two species of Artocarpus. **B.** Repeat landscapes for 14 species of order Rosales showing the distribution of different repeat types and their ages of insertions depicted by Kimura distance (along the x-axis) and Genome coverage (along y-axis) percentages. *A. heterophyllus* shows a peak (3.5% of genome coverage) of LTR-Copia abundance during the recent period (~0.05 Kimura distance), which is not shared by the other two species as they have a peak (~2.5% of genome coverage) during the comparatively older period (~0.1 Kimura distance). Apart from these, *P. persica* shows the most recent accumulation of DNA CMC repeats, which might contribute to its incipient evolution.

Similarly, *F. vesca* and *P. persica* have an unusual abundance of DNA CMC repeats but have accumulated at different Kimura distances. While the repeat content in *F. vesca* has peaked at a Kimura distance of ~0.05, the accumulation of repeats in *P. persica* appears to be more recent. Further research is required to understand if this represents an ongoing insurgence of DNA CMC by comparing high-quality genomes and transcriptomes of closely related species/varieties.

### Species-specific signatures of selection

Genes identified as positively selected by the three approaches (i.e., PAML, aBSREL, and RELAX) were considered to reduce false positives. While this approach identifies a smaller set of genes, these results are more reliable and robust to the approaches employed (see **Supplementary Figure 4**). We discuss the pathways with several positively selected genes in a comparative framework to understand putative species-specific adaptations.

#### Starch and sucrose pathway (see **Figure 7A**)

Starch and sucrose metabolism is at the heart of plant growth and development. All three Artocarpus species shared signatures of positive selection in genes producing, (1) GBE1 (1,4 alpha-glucan branching enzyme) involved in the Starch synthesis step and (2) Cellulase/endoglucanase involved in the breakdown of cellulose. The genes coding for BGLU (Beta-Glucosidase) and EGLC (Glucan endo-1,3-beta-D-glucosidase) are involved in synthesizing D-glucose, through the production of multiple intermediate metabolites, were positively selected in both *A. hirsutus* and *A. altilis*. However, there are multiple species-specific shifts in selection strength among the three Artocarpus species. For instance, *A. hirsutus* has multiple positively selected genes in different subprocesses of the starch and sucrose metabolism pathway and includes all the genes involved in the conversion of UDP-glucose to D-glucose through the production of Trehalose-6-P and alpha-Trehalose, i.e., otsA (Trehalose 6-phosphate synthase), otsB (Trehalose 6-phosphate phosphatase) and TREH (Trehalase). The BAM (Beta-amylase) gene involved in the breakdown of starch into Dextrin and Maltose through Maltodextrin was also positively selected in *A. hirsutus*. Interestingly, none of these genes had any signatures of selection in the other two Artocarpus species. Similarly, *A. altilis* has a species-specific positive selection in the PGM (Phosphoglucomutase) gene involved in converting D-glucose-1-phosphate to D-glucose-6-P. The comparative analysis of positively selected genes in this pathway highlights the differential regulation of plant developmental processes, especially in *A. hirsutus*, which has several positively selected genes in Trehalose synthesis and metabolism.

**Figure 7:**
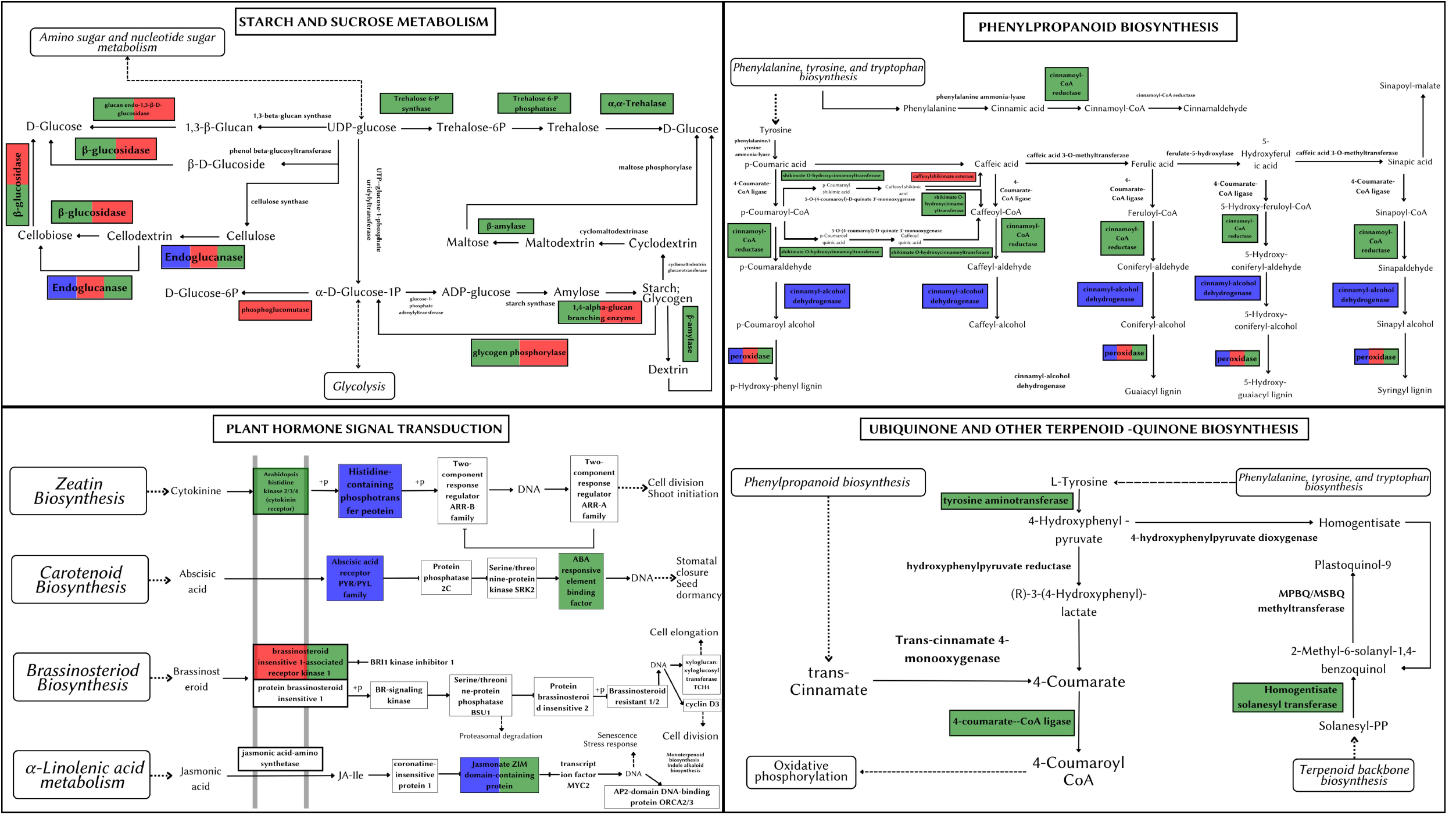
Metabolic pathways depicting positively selected genes in Artocarpus species. The positively selected genes in *A. hirsutus* (green), *A. heterophyllus* (blue), and *A. altilis* (red). **A.** Starch and Sucrose Metabolism. **B.** Phenylpropanoid biosynthesis. **C.** Plant hormone signal transduction. **D.** Ubiquinone and other-terpenoid-quinone biosynthesis.

#### Phenylpropanoid biosynthesis/Lignin pathway (see **Figure 5B**)

The phenylpropanoid pathway is involved in the biosynthesis of secondary metabolites such as lignins and flavonoids using Phenylalanine, tyrosine, and tryptophan-derived compounds. All three species of Artocarpus have signatures of positive selection in the genes producing peroxidase (PRX/PRDX) enzyme, which catalyses the last step of lignin biosynthesis by converting lignin alcohols to lignins. Species-specific positive selection is detected in *A. hirustus* among the genes involved in the pathway’s initial stages, such as 4CL (4-coumarate-coA ligase) and HCT (shikimate O-hydroxycinnamoyltransferase). Similarly, positive selection is detected in CAD (Cinnamyl-alcohol dehydrogenase) and CSE (Caffeoyl shikimate esterase) for *A. heterophyllus* and *A. altilis*, respectively. 4CL is common to both lignin and flavonoid biosynthesis pathways, while CAD, HCT, and CSE are considered lignin pathway-specific genes (Falcone Ferreyra et al., 2012; Yao et al., 2021). However, HCT is also thought to have a role in flavonoid biosynthesis (Ren et al., 2020).

#### Plant hormone signal transduction (see **Figure 7C**)

Hormone signal transduction involves numerous but crucial critical players in plant development. A transmembrane protein, BAK1 (Brassinosteroid insensitive 1-associated receptor kinase 1), is a co-receptor of BRI1 (Brassinosteroid insensitive 1) and plays a vital role in development, stress tolerance, and plant-pathogen interactions. BAK1 is positively selected in *A. hirsutus* and *A. altilis* but not in *A. heterophyllus*, suggesting differential developmental regulation in these species. Apart from BAK1, *A. hirsutus* has elevated selection pressure on another transmembrane protein HK2/3 (Histidine Kinase 2/3), the receptor for cytokinin, which is instrumental in shoot initiation and vascular bundle formation. ABF (ABA-responsive element binding factor) protein involved in regulating plant abiotic stress responses is also positively selected in *A. hirsutus*. Another important factor involved in JA (Jasmonic Acid) pathway, JAZ (Jasmonate ZIM domain-containing protein) (Pauwels and Goossens, 2011), also experiences higher selective pressure in *A. hirsutus* and *A. heterophyllus*. JA pathway is involved in almost every developmental process, including flower and root development and protection or response to multiple biotic or abiotic stress (Yang et al., 2019). Furthermore, *A. heterophyllus* also has two more genes that have elevated selection pressure, AHP (Histidine-containing phosphotransfer protein) and PYL (abscisic acid receptor PYR/PYL family) regulators of cytokinin and ABA (Abscisic acid), respectively.

#### Ubiquinone and terpenoid-quinone biosynthesis (see **Figure 7D**)

Ubiquinone and other quinone-related compounds participate in multiple growth and developmental processes and act as antioxidants to provide stress tolerance (Liu and Lu, 2016). Genes involved in this pathway such as 4CL, TAT (Tyrosine aminotransferase), HST (Homogentisate solanesyltransferase), COQ6 (Ubiquinone biosynthesis monooxygenase), and NDC1 (Demethylphylloquinone reductase) all were positively selected in *A. hirsutus*, whereas neither of the two other Artocarpus species had any positive selection in this pathway.

#### Carotenoid biosynthesis (see Figure 8A)

Carotenoid pathway aids in the development, stress response, and synthesis of carotenoid products (Shumskaya and Wurtzel, 2013). *A. hirsutus* has a positively selected gene PDS (15-cis-phytotene desaturase) involved in the production of carotenoids, specifically zeta-carotenoids and their derivatives. These carotenoids have a yellowish colour and are involved in fruit ripening (Naing et al., 2019). The CYP707A gene, which is involved in catabolising ABA and its regulation, is positively selected in *A. altilis*. ABA is involved in germination and other stress responses, which suggests CYP707A may be regulating seed development processes (Kim et al., 2020).

**Figure 8:**
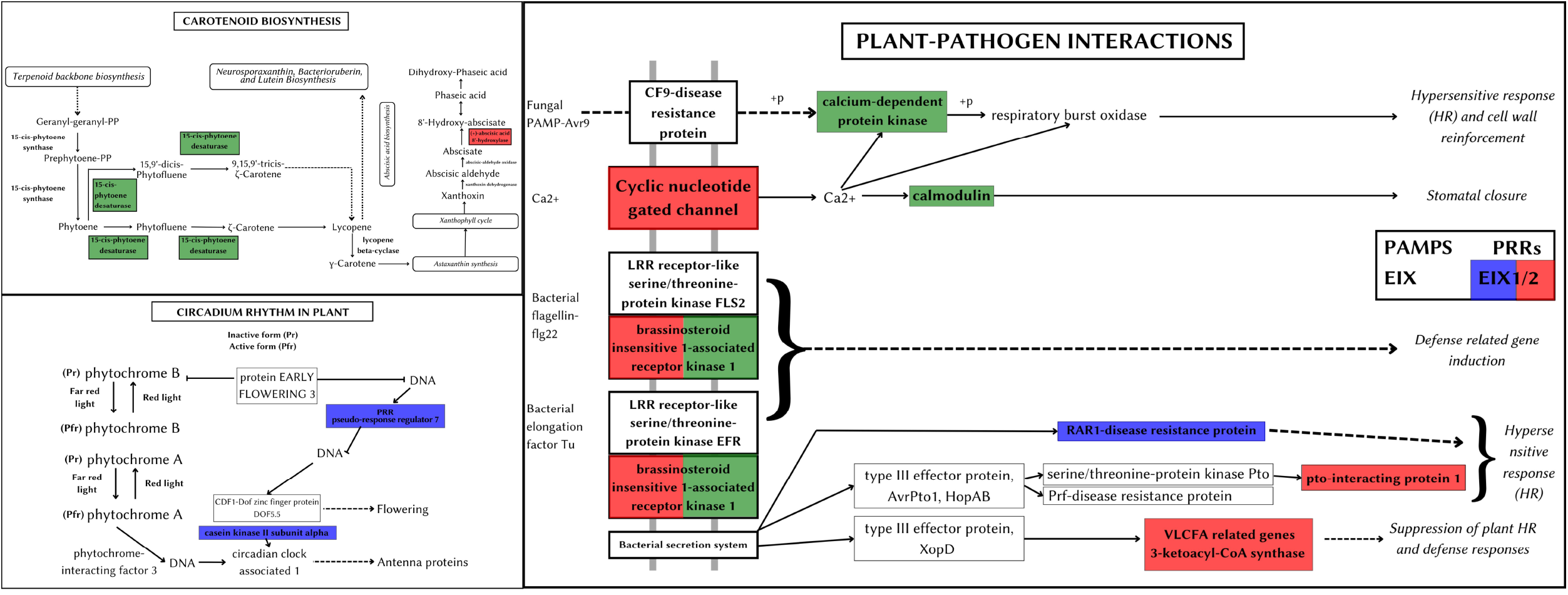
Metabolic pathways depicting positively selected genes in Artocarpus species. The positively selected genes in *A. hirsutus* (green), *A. heterophyllus* (blue), and *A. altilis* (red). **A.** Carotenoid biosynthesis. **B.** Plant-pathogen interactions. **C.** Circadian rhythm in plants.

#### Plant-pathogen interaction (see **Figure 8B**)

Plant-pathogen interactions impact the survival and development of the plant. The plant can elicit pathogen-specific immune responses by assessing the nature of the pathogen. *A. hirsutus* has three positively selected genes involved in this pathway CPK (Calcium-dependent protein kinase), CALM (Calmodulin), and BAK1. CPK and CALM are part of fungal PAMP-triggered immunity (Pathogen-Associated Molecular Pattern) and provide fungal responses. BAK1 is involved in the bacterial pathogen response of the plant. RAR1 protein (Disease resistant protein) involved in Effector-triggered immunity against bacterial pathogens is positively selected in *A. heterophyllus*. Pattern recognition receptor (PRR) EIX receptor 1/2 was positively selected in both *A. heterophyllus* and *A. altilis*. Other than this PRR, *A. altilis* showed positive selection in multiple different genes involved in plant-pathogen interactions, like CNGC (Cyclic nucleotide-gated channel), a transmembrane protein providing fungal response PTI1 (pto-interacting protein 1), BAK1 and KCS (3-ketoacyl-coA synthase) involved in bacterial defence responses.

#### Circadian rhythm in the plant (see **Figure 8C**)

Circadian rhythm of plants controls the molecular and cellular expression patterns to regulate better and mediate the light and dark periods, which in turn gives a fitness benefit to the plant (Venkat and Muneer, 2022). *A. heterophyllus* has two positively selected genes, PRR7 (pseudo-response regulator 7) and CSNK2A (Casein kinase II subunit alpha), which regulate plant circadian rhythms. The importance of efficient control of light and dark periods could also be an evolutionary adaptation that could have led *A. heterophyllus* to distribute to a larger geographical area.

#### Floral gene evolution

Floral genes AS1 (Asymmetric Leaves 1) and HUA2 (Enhancer of AG-4 2) experienced strong selection pressure in both *A. hirsutus* and *A. heterophyllus*. At the same time, TIL1 (Tilted 1), TPS1 (Trehalose-6-phosphate synthase), GA2 (GA requiring 2), and MEE27 (Maternal effect embryo arrest 27) were positively selected only in *A. hirsutus*. VIP5 (Vernalisation Independence 5) and PRR7 (Psuedo-response regulator 7) experienced elevated signatures of selection only in *A. heterophyllus*. PFT1 (Phytochrome and flowering time 1), HDA9 (Histone deacetylase 9), and LNK2 (Night light-inducible and clock-regulated 2) have lineage-specific signatures of selection in *A. altilis*. Similarly, PHYE (Phytochrome E) was positively selected in both *A. altilis* and *A. heterophyllus*.

All genes under strong signatures of selection in *A. altilis* and some in *A. heterophyllus* either had functions in light signalling and/or circadian rhythm regulation, which might be the case because of the dependence on the tight regulation of flowering time for these plants’ fitness. However, genes under strong selection pressure in *A. hirsutus* were involved in meristematic growth, fungal defense, pollen/flower development, and identity. Such processes explain the dependence on maintaining the phenotype rather than adapting to changes in light periodicity. But, *A. heterophyllus* experiences selection pressure on both circadian and other phenotype-related genes and hence might have allowed the evolution of a distinctive phenotype with a wider distribution.

## Discussion

Our newly generated Wild Jack genome allowed comparative genomics of phenotypically diverse, phylogenetically distant breadfruit species from contrasting habitats. Changes in gene content between these species reflect putative modifications in the distinct phenotypes of these plants. Notably, several genes from the same biological processes, such as flower development and response to biotic/abiotic stress, have experienced changes in copy number and may have allowed the rewiring of these pathways. Signatures of selection also occur in some genes of these same pathways leading to fine-tuning the altered phenotypes. Whole-genome sequence data allows the reconstruction of the demographic history and comparison between closely related species. Hence, such reconstructions provide helpful insight into habitat-specific responses. We complemented the information from these genomics-based reconstructions with species distribution modeling. Broadscale patterns identified by species distribution models were concordant with the trajectory inferred by genomics-based reconstructions and indicated that Jackfruit and Wild Jack species, which share the same habitat, have a comparable population size history. In contrast, the Breadfruit, which has a native range with a history of volcanic eruptions, has a unique history of population bottlenecks. While the effect of habitat on the demographic history is expected, the response of each Artocarpus species to the habitat can be idiosyncratic.

### How has the habitat shaped the genomes of Artocarpus?

The demographic reconstruction and species distribution modelling revealed the effects of differentiated bioclimatic forces acting on their bottleneck history and distribution patterns in two diversified habitats, i.e., the Western Ghats (Jackfruit and Wild Jack) and East of Sulewasi (Breadfruit). The oceanic region of East Sulewasi islands has a history of volcanic eruptions, with accumulated ash resulting in acidic soils. These acidic soils have unique properties and minerals or microelements. The climate and precipitation cycles are also different compared to the Western Ghats. In contrast, the Western Ghats have nutrient-rich, alkaline soils with abundant biotic meta compositions of various taxa in the soil (Myers et al., 2000). This nutrient-rich soil contains numerous bacterial and fungal pathogens, and plants must adapt to achieve fitness in interacting or responding to these species. All these differences have impacted the flora and fauna of these regions, and hence these plants must adapt to differential plant-biotic interactions. Breadfruit (*A. altilis*) has multiple gene family expansions in the OXPHOS assembly factor and chloroplast genes. We also identified lineage-specific copies of the organellar expression regulator transcription factor mTERF15 and acidic soil response transporter ABCB27. These multiple expansions in energy-producing pathways can be explained by the higher energy demand of the plant to maintain oxidative stress homeostasis and acquire resistance to Aluminium-rich acidic soils. Due to harsh soil and bioclimatic properties, Breadfruit, mitochondrial assembly genes, and their regulators have experienced gene family expansion.

Jackfruit and Wild Jack have multiple gene family changes related to plant-biotic interactions and secondary metabolite productions, which are important determinants for biotic and abiotic adaptations. The Wild Jack shows gene family expansions for IRE1, GRIP, SCPL-II, PTR1, DTX28, HSP20, MDL3, and receptor kinases, all involved in either biotic, abiotic stress tolerance/response, or plant immunity. Similarly, the Jackfruit has gene family expansion in the stress-related GST gene family and unique gene clusters of genes like the TIR domain gene involved in plant immunity and the RLP gene family involved in stress responses. In addition, both Western ghat species have signatures of selection in anti-fungal genes such as AS1, DMS11 (Defective in meristem silencing 11), RD20 (Responsive to desiccation 20), LECRK-IX.1 (L-TYPE LECTIN RECEPTOR KINASE IX. 1) conferring fungal-resistant properties to its timber.

### Why such divergent phenotypes among Artocarpus trees?

Animals treasure the fruits of Jackfruit and Wild jack as they are sweet, fleshy, and nectary. However, the Breadfruit is a starchy fruit that is not as sweet and nectary as the other two; hence it is eaten as a vegetable rather than fruit. As discussed above, due to higher energy expenditure to maintain oxidative stress homeostasis, many other pathways with relatively more minor functions might have been impacted, reduced, or relaxed and could explain the loss of sweet and nectary fruit phenotype in Breadfruit. Even the two Western Ghats species, Jackfruit and Wild Jack, differ in their plant height, branching, fruit size, colour, etc. Although the distribution of these plants overlaps, their responses to similar biotic environments may have been different due to their contrasting growth patterns. Wild jack is a typical forest-adapted species with large trees having unidirectional growth, maintaining the apical branching to compete for sunlight efficiently, and a strong tap root system to utilize water and nutrients in dense forests. Due to this phenotype of Wild Jack, the lineage-specific gene family expansions, unique gene clusters, and genes showing selection signatures are primarily attributed to plant-pathogen interactions, stress responses, and floral evolution.

The fruits of Wild jack are at a greater height, reducing its niche of vertebrate land dispersers such as elephants, boars, and other ruminants. The increased height ensures a different mechanism for both pollination and dispersal. The long and stalky male inflorescences of Wild Jack, in contrast to the short cylindrical inflorescences of Jackfruit, might be a switch from faunal dependence for pollination to a wind-pollinated mechanism and can explain multiple gene family expansions, unique gene clusters, and positive selection in pollen recognition genes from the lectin gene family, the Receptor kinases. Due to the switch to wind pollination, the plant must have devised some mechanisms to maintain Self Incompatibility (SI). The receptor kinases are well known to function in maintaining SI to avoid self-pollination and allow cross-pollination as much as possible (Sherman-Broyles et al., 2007). The number of fruits is more and has distinctively bright yellowish-orange colour and smaller sizes as compared to others which is an adaptation for attracting the birds, bats, and primates as their dispersers (Primack, 2003; Flörchinger et al., 2010). The Lion-tailed Macaque (*Macaca silenus*), endemic to the Western Ghats, is one of the most important consumers of these fruits and can be considered their dispersers (Kumara and Santhosh, 2013). Some hornbills have also been observed eating these fruits. These pollination/disperser-specific changes in Wild Jack might be due to gene family changes and changes in selection pressure in floral genes like AGL15/18, ARA1F, PTR3, EDA17, RK3, TIL1, TPS1, GA2, MEE27, AS1, HUA2, etc. In comparison, the colour of fruit could be due to strong selection pressure on carotenoid biosynthesis genes like PDS, which is crucial for synthesizing zeta-carotene, which has a yellowish pigment. All these genomic changes have translated into the phenotype of the Wild Jack to adapt to the forest habitat and fine-tune its pollination and disperser network.

Trehalose metabolism contributes to processes involving embryogenesis, flower and plant development, branching of inflorescence, etc. (Lunn et al., 2014). Additionally, TPS1 regulates axillary bud outgrowth and modulation of axial shoot branching (Fichtner et al., 2021). In the Wild Jack genome, all the trehalose metabolism genes are positively selected, suggesting its importance in maintaining the phenotype of apical branching and changes in inflorescence structure. Moreover, Wild Jack has a gene family expansion in F6’H (Feruloyl-CoA 6’-hydroxylase) which catalyses the penultimate step in scopoletin synthesis, a simple coumarin. A recent study (Hoengenaert et al., 2022) demonstrated that elevated expression of Scopoletin in lignifying cells leads to higher production of monosaccharides. Due to the higher lignocellulosic mass of Wild Jack, the Trehalose pathway’s involvement in generating sugars and their conduction seems likely in this plant.

In contrast to Wild Jack, Jackfruit has a short, branched tree structure with low-hanging fruits that are not suited for dense forests. The large fruits of Jackfruit are nectary and sweet with inflorescences which also impart volatile compounds which attract a species of Gall Midge that may facilitate pollination (Gardner et al., 2018). The low-hanging Jackfruit is consumed by large mammals like elephants, wild boar, and other ruminants, facilitating its dispersal. Long-range effective dispersal is possible due to the long-distance migration of these dispersers. These unique phenotypes of Jackfruit allow efficient fauna-based pollination/dispersal mechanisms. Gene family expansions and lineage-specific selection among genes of the flavonoid biosynthesis pathway, like Chalcone-synthase, could have facilitated the evolution of nectary fruits and inflorescences with volatile compounds. The widespread distribution of Jackfruit spans regions with differing light periodicity and would need to adapt to these changes. The strong signatures of selection in the genes involved in light signaling or circadian rhythm suggest a tight regulation of light periodicity-related pathways. An efficient plant-pollinator/disperser network and tight regulation of circadian rhythm might have played an instrumental role in maintaining the wider distribution range and larger population size.

### Did LTR-Copia accumulation shape Jackfruit evolution?

We observed recurrent genome size changes due to repeat content dynamics in Rosales’ order. We also see that the genomes of order Rosales show a strong positive correlation between their genome sizes and repeat content (**Figure 5B**). The increase in genome size with repeat content suggests that genome size evolution is influenced by repeat expansion. The size of the assembled *A. heterophyllus* (Jack fruit) genome (~980 Mb) is ~200Mb larger than that of *A. altilis* (~800 Mb) and *A. hirsutus* (~790 Mb). Assembled genome sizes concord with the K-mer-based estimates and is largely unaffected by assembly quality. Moreover, the genomes of *A. heterophyllus* and *A. altilis* are from a single study (Sahu et al., 2019) that uses the same methodology for sequencing and assembling both genomes, ensuring comparable genome quality. The difference in genome size among the Artocarpus species is primarily due to the increased prevalence (~150 Mb) of LTR-Copia in *A. heterophyllus* (*see* **Figure 5A**). The genome of *C. sativa* from the sister family Cannabaceae also has a larger genome, potentially due to a lineage-specific accumulation of LTR-Copia. Investigation of repeat accumulation dynamics suggests recent lineage-specific repeat expansions in these two phylogenetically distant species in a similar time frame, which suggests a role of habitat or stress-mediated induction of repeats. Repeat content change in plants has been linked to functional diversification through cis-regulatory changes or other epigenetic mechanisms (Negi et al., 2016; Hirsch and Springer, 2017). The conflict between transposable elements and the host defense mechanisms is elevated in stress conditions resulting in improved regulatory machinery (Wang et al., 2018). Hence, the accumulation of LTR-Copia in *A. heterophyllus* has played a pertinent role in the evolution of the Jackfruit genome. Gene expression data will allow the identification of ongoing transposon activity and its effect on gene regulation.

### Limitations and broader implications

Fragmented genomes lead to gene content underestimation, and genome quality heterogeneity can confound comparative genomics. All three Artocarpus genomes compared in this study have similar BUSCO scores and are fairly comparable in gene content. Additionally, we put forth multiple stringent criteria to avoid false positives. We identify several candidate pathways that have experienced changes in gene content and positive selection. Detailed functional characterisation of these candidates by evaluating changes in gene expression and the consequent phenotypic changes will require further studies. The occurrence data for Artocarpus is limited and influenced by human-mediated dispersal, which could confound the species distribution modelling. The single genome-based demographic history reconstructions performed using the PSMC method are known to be unreliable in the recent past (<20KYA). Future studies can provide better resolution by incorporating population-level sampling.

Of the ~70 species of Artocarpus, our study includes only three whole genomes. Although our study highlights the potential of such comparative genomic studies, the inclusion of multiple other species would be able to provide definitive answers to questions regarding the origin, phenotypic diversity, and diversification. For instance, the genetic basis of syncarp evolution in this genus can be explored to exploit the molecular mechanisms involved in achieving desired phenotypes. Such species-rich genera with heterogeneous phenotypes are especially well suited for agroforestry genomics. Hence, Artocarpus can serve as a model to understand inflorescence/syncarp biology.

## Conclusion

Our study has generated genomic resources for a forest tree, the Wild Jack, which is endemic to the Western Ghats. This dataset will help understand the evolution of forests and fill a gap in sampling forest tree genomes. Comparative genomic analysis with other Artocarpus species and members of the order Rosales has provided interesting insights into their genomic evolution. Habitat-driven evolution through phenotypic diversification has resulted in genomic signatures of selection and gene-family changes. Demographic history reconstructions from genomic data and species distribution modelling strongly support the prominent role of habitat. Adaptive changes in plant growth and development, floral morphology, and biotic interactions have shaped the Wild Jack to thrive in the forests and may explain its endemism and current fragmented distribution. In contrast, Jackfruit and Breadfruit appear tightly regulated by light signalling and circadian rhythm leading to more widespread distribution. Additionally, the fruit morphology/sizes might be due to genic evolution in floral development and may be due to the habitat-specific rewiring of the pollinator/dispersal network.

## Acknowledgment

We thank the Ministry of Human Resource Development fellowship to ABP. The Department of Biotechnology, Ministry of Science and Technology, India (Grant no. BT/11/IYBA/2018/03) and Science and Engineering Research Board (Grant no. ECR/2017/001430) provided funds used to generate primary sequencing data published in this article and computational resources (i.e., Har Gobind Khorana Computational Biology cluster) used. We want to thank the lab members of the PCDB lab, IISER Bhopal, for their valuable discussions.

## Author contributions

A.B.P and N.V. wrote the manuscript with inputs from S.S.V and C.G.K. S.B.N, and R.S collected the samples required for primary data generation. A.B.P analyzed the genomic data and generated all the results. Species distribution modeling analysis was done by S.S.V, who also prepared the illustrations used in this manuscript. All authors reviewed the manuscript.

## Availability of data

The primary sequencing data are available from the ENA under BioProject# PRJEB55580. All **Supplementary information**, including other associated data and scripts used for analysis, are provided in an easy-to-browse format: https://github.com/Ajinkya-IISERB/Wild_Jack.

## Contribution to field

The Western Ghats is a biodiversity hotspot with high levels of endemism. Recent developments have threatened the population of many endemic species in this region. Hence, quantifying the diversity and distinctiveness of species from the Western Ghats is a pressing concern. However, a scarcity of genomic datasets, especially from forest trees, impedes prioritizing conservation policies. Due to the endemism and the lack of natural history information about these species, their commercial value is underutilized and tends to get neglected in conservation efforts. We focused on a multi-purpose endemic forest tree with closely related commercially successful species with genomic resources to highlight the importance of conserving such forest tree species. In this study, we sequence, assemble and annotate the genome of the Wild Jack (*Artocarpus hirsutus*) sampled from the sacred groves of the Western Ghats. Comparing the Wild Jack genome with the commonly cultivated species *Artocarpus heterophyllus* (Jackfruit) and *Artocarpus altilis* (Breadfruit) provides important clues about the genetic distinctiveness and phenotypic diversification in the Artocarpus genus. We identify several candidate genes from multiple pathways, which can be the focus of further research into their phenotypic implications that functional studies can explore.

## Supplementary Figure legends

**Supplementary Figure 1: Chloroplast genome of *A. hirsutus* and rearrangements in Genus Artocarpus.**

A. The assembled chloroplast genome of *Artocarpus hirsutus* with a length of ~161 Kbp.

B. Whole plastome-based bootstrapped phylogeny of Artocarpus and outgroup species *Ficus religiosa* and *Morus indica*. The second panel shows Mauve alignment of variable regions of these chloroplast genomes. All Artocarpus genomes show a similar arrangement of inverted repeat and small single copy (SSC) regions except *A. heterophyllus* and *A. integer*. These two genomes show an inverted arrangement of sequences and genes of the SSC region.

**Supplementary Figure 2: The phylogenetic relationship between gene copies of the Lectin gene family Receptor Kinase 3 (RK3).**

Gene copies of *Artocarpus hirsutus* are coloured in green, *A. heterophyllus* in blue, and *A. altilis* in red.

**Supplementary Figure 3: Phylogenetic relationship between gene copies of ASMT and COMT gene families.**

The sky blue coloured cluster of genes is the COMT gene family, whereas the red coloured is ASMT. Gene copies of *Artocarpus hirsutus* are coloured in green, *A. heterophyllus* in blue, and *A. altilis* in red.

**Supplementary Figure 4: Approaches used to detect positive and relaxed selection in Artocarpus species.**

Genes identified as positively selected by all the approaches are shortlisted as positively selected genes.

