## Supplementary figures and images for "Jack of all trades: genome assembly of Wild Jack and comparative genomics of Artocarpus"

### Supplementary Figures

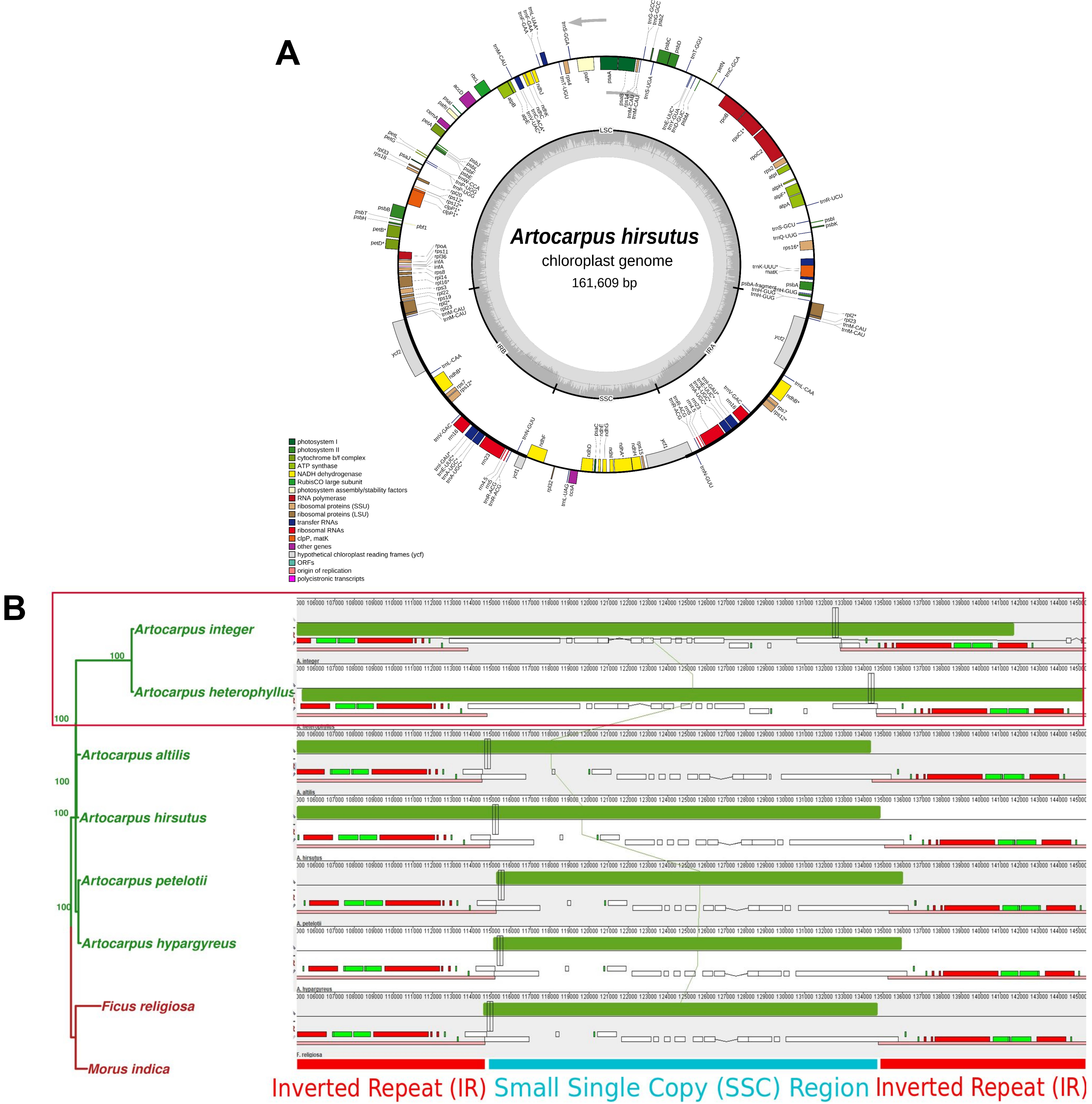

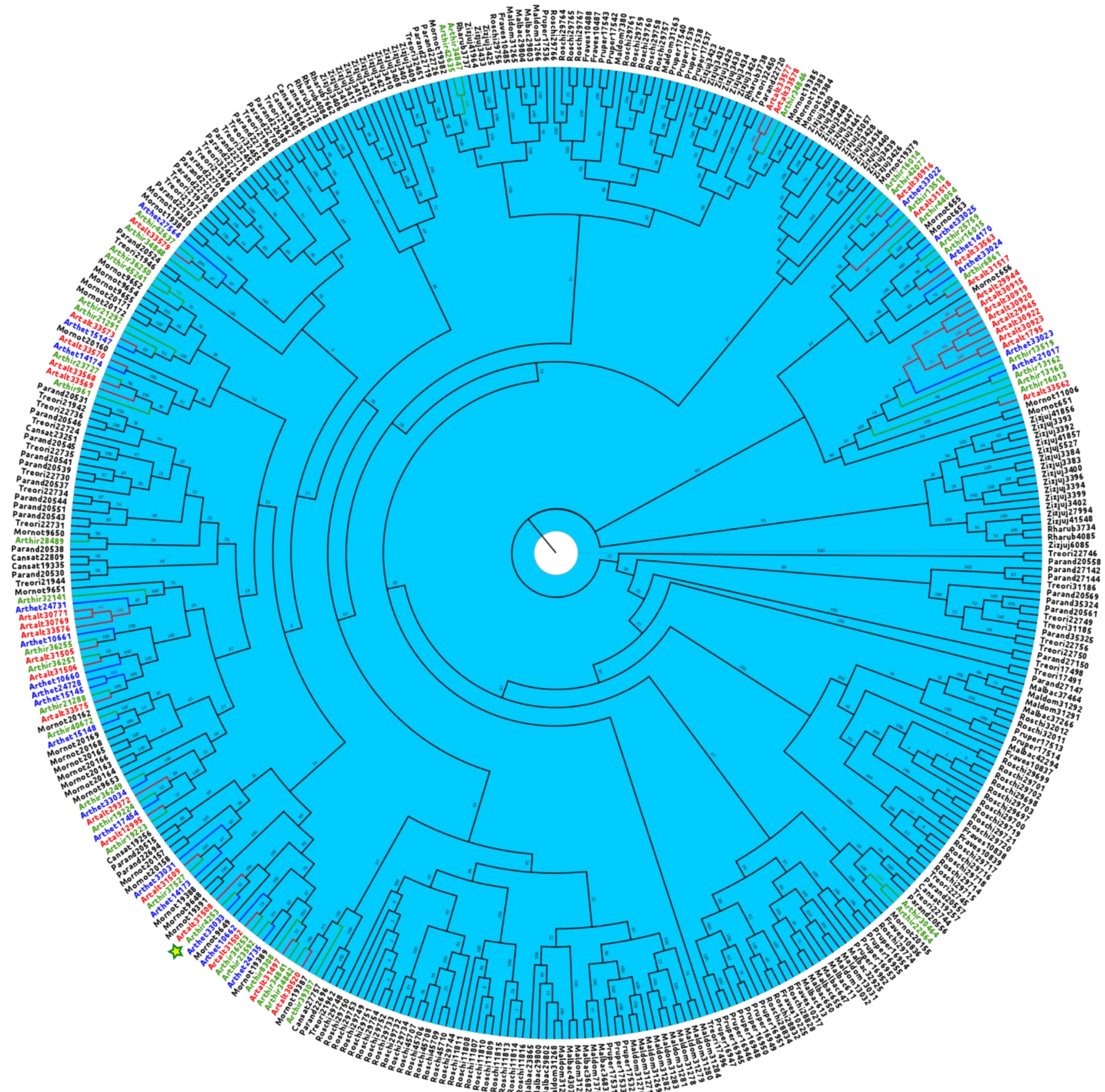

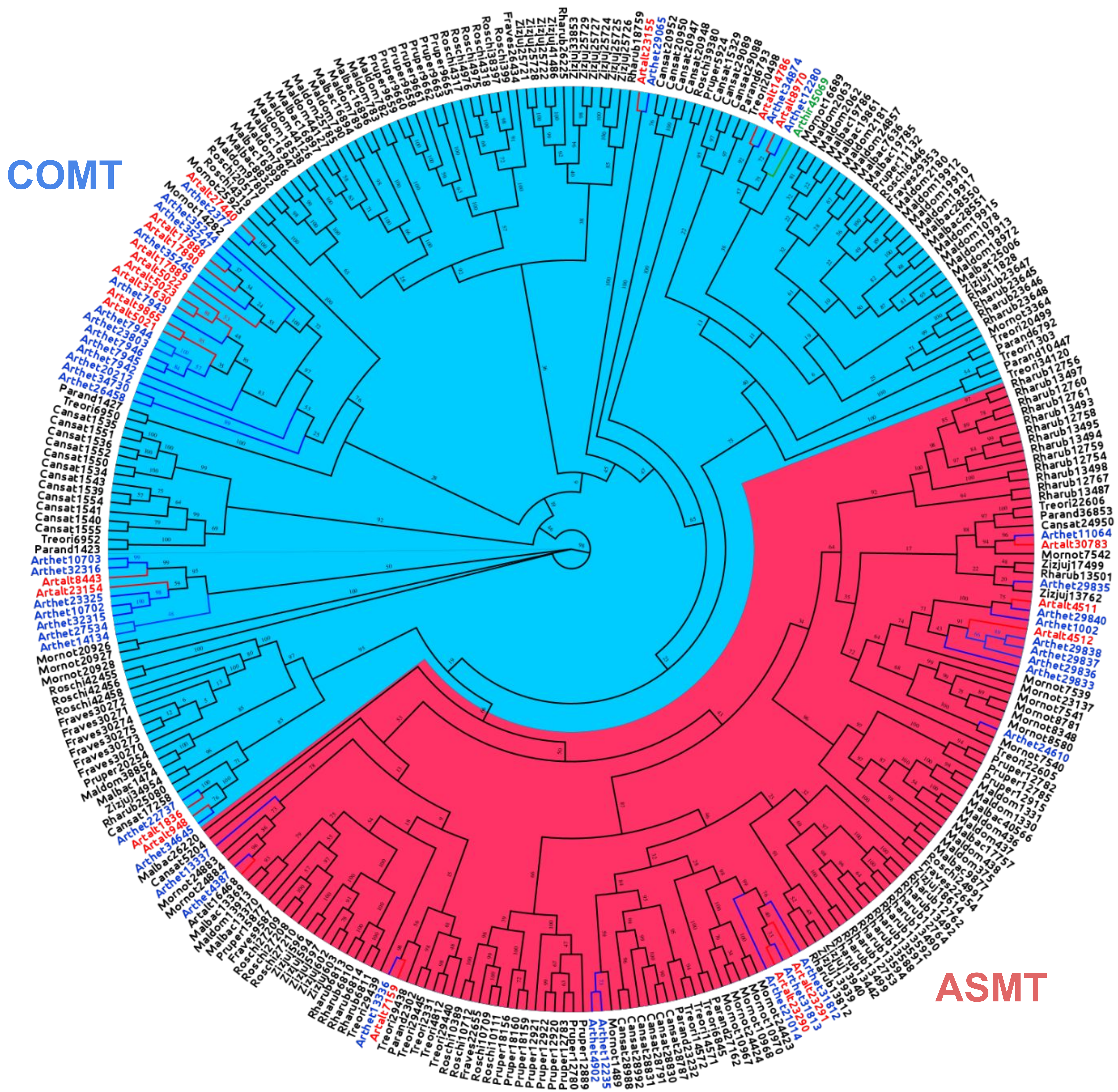

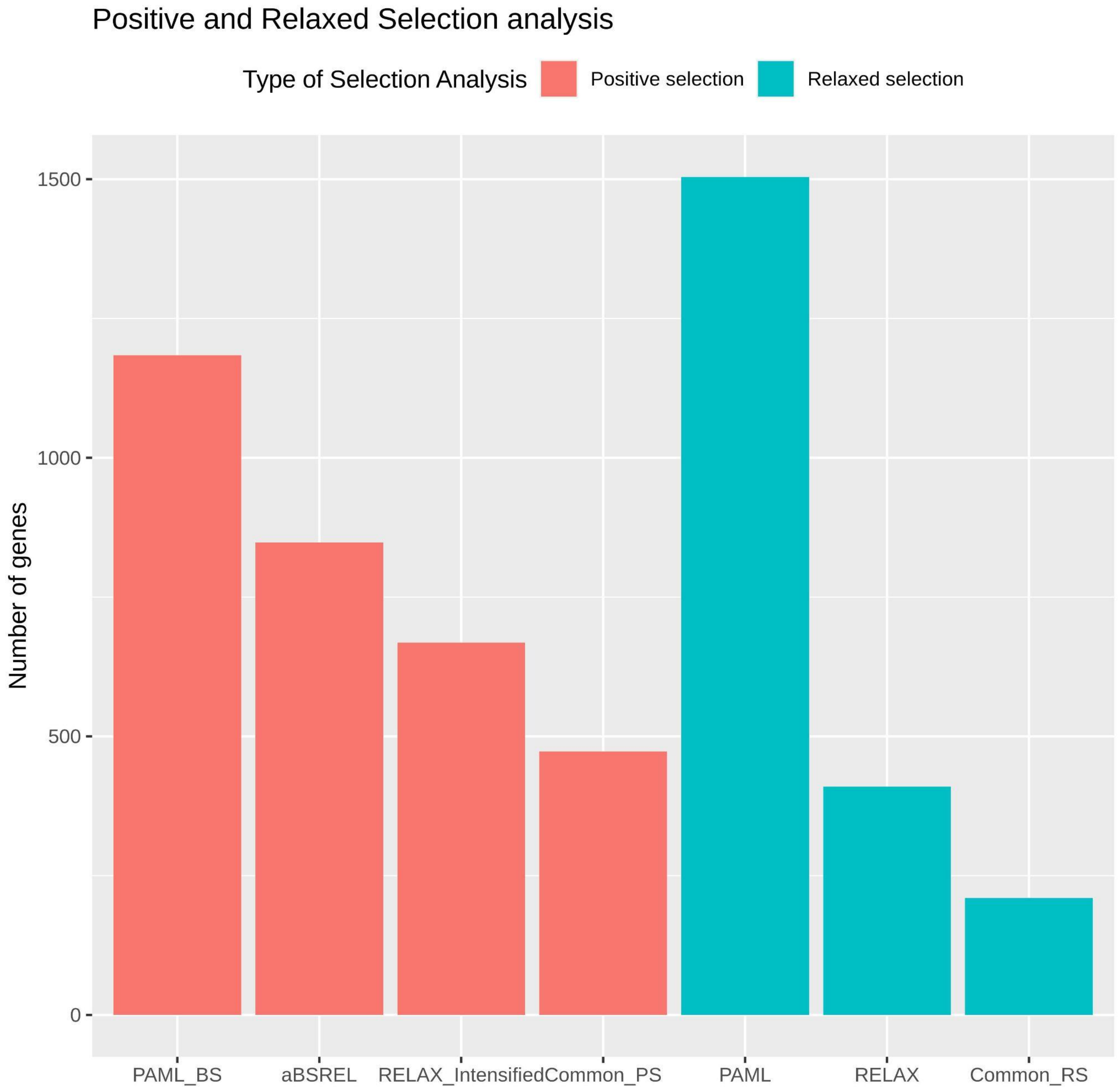
